# Development of a CRISPR/Cas9-mediated transformation procedure for the wheat pathogen *Zymoseptoria tritici*

**DOI:** 10.64898/2026.05.27.728285

**Authors:** Sandra V. Gomez-Gutierrrez, Maikel B. F. Steentjes, Gert H. J. Kema, Stephen B. Goodwin

## Abstract

*Zymoseptoria tritici* is the causal agent of Septoria tritici blotch (STB), one of the most destructive diseases of wheat worldwide. Although the *Z. tritici* genome encodes hundreds of predicted effector proteins, functional characterization through the use of genome-editing techniques has been limited due to low homologous recombination efficiency and extensive effector redundancy. In this study, we established and evaluated a CRISPR/Cas9-based genome editing procedure for targeted effector gene disruption in *Z. tritici* using in vitro–assembled Cas9–sgRNA ribonucleoprotein (RNP) complexes combined with short (60 bp) homologous donor DNA flanks. Using this approach, we successfully generated knockout mutants for a selected candidate effector gene, the Hce2 domain–containing effector Mycgr3107904.

Virulence assays on the susceptible wheat cultivar Taichung 29 revealed that two independent *ΔMycgr3107904* mutants exhibited a pronounced delay in symptom development compared to the wild-type strain IPO323, with disease onset and progression delayed by approximately 4–5 days. While mutant strains ultimately followed a similar disease trajectory, wild-type–infected leaves displayed extensive necrosis and pycnidia formation at earlier time points, indicating a significant reduction in virulence upon loss of Mycgr3107904. Together, our results demonstrate the feasibility of CRISPR/Cas9-mediated effector gene knockout in *Z. tritici* and provide functional evidence that Mycgr3107904 contributes to timely disease progression. This work advances genome editing tools for *Z. tritici* and facilitates systematic dissection of effector functions underlying fungal virulence.

## Background

*Zymoseptoria tritici* is a major fungal pathogen of wheat and the causal agent of Septoria tritici blotch (STB). It is considered one of the most persistent and economically damaging threats to wheat production worldwide, and affects both bread and durum wheat, particularly in temperate regions [1]. In fields planted with STB-susceptible wheat cultivars, severe epidemics can result in yield losses of up to 50% [2]. STB stands as the most prevalent foliar disease in Western Europe and remains a major concern in global wheat cultivation [3]. Given the substantial impact of *Z. tritici* on wheat production, extensive research has focused on understanding the molecular interactions with its host. These efforts have led to the identification and functional characterization of several effector proteins [4–11].

The development of transformation techniques to generate *Z. tritici* knockout strains with disruptions of targeted effector genes has significantly enhanced the ability to investigate effector functions, particularly in pathogen recognition and host immune evasion [12]. Key advancements include PEG-mediated transformation protocols [13], vectors with various selectable markers [12], *Agrobacterium tumefaciens-*mediated transformation [7,14–16], GFP reporter constructs [17], gene over-expression [18] and high-throughput techniques for rapid functional screening in *Z. tritici* [19]. Despite these many advances in genetic manipulation techniques for *Z. tritici*, challenges persist. The fungus exhibits a low frequency of homologous recombination [14], complicating homology-mediated targeted gene disruptions. These challenges are further exacerbated by the functional redundancy among *Z. tritici* effectors [20], making phenotypic effects difficult to detect following single-effector gene knockouts . To address these issues, advanced gene-editing technologies, such as CRISPR/Cas9, offer more efficient strategies for precise genetic modifications in *Z. tritici*.

The clustered regularly interspaced short palindromic repeat (CRISPR) and CRISPR-associated protein 9 (Cas9), collectively known as CRISPR/Cas9, is a groundbreaking genome-editing technology originally identified as an adaptive immune system in bacteria and archaea, protecting them from invasive nucleic acids [21–23]. CRISPR/Cas9 has been applied for gene knockout, insertion, and replacement [24]. CRISPR/Cas systems are categorized into two classes based on the type of Cas proteins they employ. The Type II CRISPR/Cas9 system from *Streptococcus pyogenes* is the most extensively studied and utilized due to its simplicity, specificity, and efficiency [25]. This system requires only two components for gene targeting and cleavage: the Cas9 endonuclease and a single guide RNA (sgRNA) that is complementary to the target DNA [23,26]. The sgRNA consists of a 20-nucleotide sequence named the protospacer, which is complementary to the target sequence. These 20 nucleotides are fused to approximately 80 nucleotides required for Cas9 binding [25,26]. The Cas9-sgRNA complex recognizes, binds, and cleaves double-stranded DNA sequences that match the protospacer, provided they are flanked by a protospacer-adjacent motif (PAM), resulting in a double-strand break (DSB) in the DNA [21,26].

Once the Cas9 nuclease induces a DSB at the target site, it can be repaired by either the error-prone, non-homologous end-joining (NHEJ) pathway or the more precise, homology-directed repair (HDR) mechanism. HDR requires the presence of a donor DNA template with flanking sequences matching the break site or the target gene, allowing precise modifications [27,28]. When coupled with an in vitro-assembled Cas9 ribonucleoprotein (RNP) complex, HDR is highly effective for complete gene knockouts [24]. However, in some fungi, such as *Aspergillus niger* and *Penicillium digitatum*, HDR events occur infrequently, with NHEJ serving as the dominant DNA repair mechanism [29,30]. In fact, homologous recombination events in *Z. tritici* occur at highly variable but generally low frequencies, ranging from 0.5% to 75% during homology-directed repair events for gene disruption [14].

The CRISPR/Cas9 system has rapidly advanced as a powerful genome-editing tool in filamentous fungi [31], offering a solution to overcome their low frequencies of homologous recombination. CRISPR/Cas9-mediated transformation has been developed in many fungal plant pathogens including the basidiomycete *Ustilago maydis*, and the ascomycetes *Botrytis cinerea* [32], *Pseudocercospora fijiensis* [33]*, Magnaporthe oryzae* [32], *Bipolaris sorokiniana* [34], and *Fusarium graminearum* [35]. In *Z. tritici,* to our knowledge there is only one study that attempted to implement CRISPR/Cas9 for the knock-out of the *NiaD* (nitrate reductase) and *PyrG* (orotidine 5-phosphate decarboxylase) genes [36]Khan et al. [36] initially employed a *Z*. *tritici* strain expressing Cas9 and introduced sgRNA into protoplasts via PEG-mediated transformation. While 14 colonies emerged on selective media, sequencing confirmed the absence of mutations in *PyrG*. To enhance transformation efficiency, they attempted direct delivery of a pre-assembled Cas9-sgRNA ribonucleoprotein (RNP) complex, but this also failed to induce mutations. To determine whether the lack of mutations was gene specific, a different gene, *NiaD* (nitrate reductase), was targeted by combining the RNP complex with a donor template designed for HDR. This donor included a hygromycin-B resistance gene flanked by 1-kb sequences matching the *NiaD* cleavage site. Although 17 colonies were recovered on selective media, none displayed successful integration of the donor cassette [36].

To better understand the role of individual effectors in *Z. tritici* virulence, we focused on generating knockout strains for selected candidate effector genes [37]. The *Z. tritici* genome is predicted to encode approximately 370 effector genes, yet only around 56 have been functionally characterized to date [38]. Additionally, transcriptome analyses in *Z. tritici* during different stages of infection and types of interaction [9,37,39–41], along with functional characterization of candidate effectors [42,43] have identified several effector genes with potential roles in virulence. Generating knockout strains for promising effector candidates is essential for testing their functional significance in *Z. tritici* virulence.

In this study, we used a CRISPR/Cas9 system with an *in-vitro* assembled RNP complex (Cas9 and sgRNA) and a donor DNA template with short homology flanks (60 bp) to facilitate microhomology-directed repair (MDR) that were injected into the host protoplasts through PEG-mediated transformation, followed by the integration of the donor DNA template at the Cas9 cutting site. With this approach we aimed to test whether CRISPR/Cas9 could be effectively used to knock out candidate effector genes in *Z. tritici.* We introduced two main modifications in this approach. First, the use of short homologous flanks of 60 bp in the donor DNA template instead of the 1-kb homologous DNA flanking the cut site of the target gene; and second, the injection of the RNP complex into the protoplasts instead of expression of the Cas9 protein. Using this novel approach, we obtained two transformant colonies. Our findings contribute to the development of an efficient genome-editing tool for generating knockout mutants, demonstrating its effectiveness for disrupting candidate effector genes, an area of extensive research in *Z. tritici*.

## MATERIALS AND METHODS

### *Z. tritici* strains and growth conditions

All experiments were done with the wild-type *Z. tritici* reference strain IPO323, originally isolated from the Netherlands [44] and reported to contain all 21 chromosomes, including 13 core and 8 accessory chromosomes in the first complete genome assembly generated for this pathogen [45]. To determine the culture medium that optimizes blastospore production for protoplast generation, the strain was cultured on potato dextrose agar (PDA), yeast sucrose agar (YSA), and malt sucrose agar (MSA). The IPO323 strain used in this study was maintained in the Laboratory of Phytopathology at Wageningen University and Research. To obtain fungal biomass for protoplasting, IPO323 was grown initially on PDA at 18 °C for four days to obtain fresh colonies. Liquid cultures were then prepared by scraping colony surfaces with an inoculation loop into Erlenmeyer flasks containing 100 mL of potato dextrose broth (PDB) and incubating for 24 hours at 18 °C with constant shaking at 200 rpm. Spore concentrations were determined using a hemocytometer to achieve a final density of 6.0 x 10⁷ conidia/mL. Knockout strains of *Z. tritici* (ΔMycgr3107904_1, ΔMycgr3107904_2, ΔMycgr394290, and ΔMycgr3109710) and the wild-type strain were cultured on PDA supplemented with 100 µg/mL of hygromycin B (Gibco™, Thermo-Fisher Scientific, Waltham, MA, USA).

### Formation of protoplasts

Protoplasts were isolated from *Z. tritici* blastospores following protocols modified from [46], [47], and Steentjes et al. (2025) (unpublished). A 1-day-old liquid culture of *Z. tritici* IPO323, grown in PDB at 18 °C, was used as the starting material for enzymatic digestion, with a final concentration of 6.0 x 10^7^ conidia/mL. A 100-mL suspension of *Z. tritici* blastospores was transferred to two 50-mL Falcon tubes and centrifuged at 3,000 rpm for 10 minutes at 4 °C. The supernatant was discarded, and the pellet was resuspended in 50 mL of sterile water with gentle shaking. The suspension was centrifuged at 3,000 rpm for 10 minutes, and the supernatant was discarded. The pellet weight was measured, and the conidia were resuspended in a 1.0 M sorbitol solution to a final concentration of 0.2 g/mL.

Simultaneously, an enzymatic digestion cocktail was prepared by dissolving 32 mg of Yatalase (Takara, Osaka, Japan) and 100 mg of Driselase (Sigma, D9515) in 10 mL of enzyme buffer containing 0.04 g of NaH₂PO₄ and 20 mL of 1.2 M MgSO₄·7H₂O (pH 5.8). The final volume was adjusted to 25 mL with additional 1.2 M MgSO₄·7H₂O. After complete dissolution, the solution was sterilized using a 0.45-µm filtration unit, followed by addition of 60 µL of Zymolyase (#E1005, Zymo Research). The enzyme mixture was placed on a rolling bench for 30 minutes to ensure proper dissolution and homogenization.

The 1.0-M sorbitol solution containing the pellet was transferred to a new 50-mL tube, ensuring approximately 2.5 g of fresh weight (∼12.5 mL of suspension) was included. The solution was then centrifuged at 3,000 rpm for 10 minutes, and the supernatant was discarded. If the pellet weighed less than 2.5 g, multiple pellets were combined [33]. Next, 10 mL of the enzymatic digestion cocktail was added to each tube, and the pellet was resuspended by gentle inversion. The suspension was incubated on a 3D rotating mixer in darkness at 28 °C for 1 hour and 30 minutes. Following digestion, 4 mL of 600 mM sorbitol + 10 mM Tris (pH 7.5) was carefully added by pipetting along the tube wall. The tubes were then centrifuged at 3,000 rpm for 10 minutes at 4 °C. Two layers formed, with protoplasts in the upper layer and debris in the lower layer. The upper layer was collected using a 1,000-µL pipette with a cut tip and transferred to a new 50-mL tube containing 10 mL of sterile TMS buffer (91 g of D-sorbitol and 1,045 mg of MOPS, pH 6.3, in 500 mL of Milli-Q H₂O). An additional 10 mL of TMS buffer was added, and the suspension was centrifuged at 1,500 rpm for 5 minutes at 4 °C. The supernatant was discarded, and the pellet was resuspended in 5 mL of cold TMS buffer by gentle shaking. A 10-µL aliquot was taken to estimate protoplast concentration using a hemocytometer.

Protoplasts were regenerated to assess their viability in forming new colonies on regeneration medium (RM) containing 0.5 g of yeast extract, 0.5 g of casein hydrolysate, 137.5 g of sucrose, and 5 g of agarose in 500 mL of Milli-Q H₂O [33]. The medium was autoclaved at 121 °C for 20 minutes. To determine the optimal hygromycin concentration for selecting transformed colonies, RM was prepared with 25, 50, and 100 µg/mL of HygB. Before transformation, RM was maintained at 60 °C to remain in a liquid state.

### Design of sgRNA and *in vitro* synthesis

The single guide RNAs (sgRNAs) for the deletion of five candidate effector genes were designed using the web-based tool CRISPick from the BROAD Institute (https://portals.broadinstitute.org/gppx/crispick/public) following the protocol for sgRNA synthesis design by [32]. Annotated sequences of each candidate effector from the *Z. tritici* IPO 323 reference strain were used as input: Mycgr394290 (XP_003851266.1); Mycgr3107904 (XP_003855719.1); Mycgr3106502 (XP_003847739.1); Mycgr3109710 (XP_003851988.1); and Mycgr3103393 (XP_003855099.1). The following parameters were used to design the protospacer sequences: (i) reference genome: Human GRCh38; (ii) mechanism: CRISPRko; and (iii) enzyme: *SpyoCas9* (Hsu et al., 2013), which recognizes the NGG protospacer adjacent motif (PAM). Two sgRNAs were generated per target gene, selected based on the highest on-target score for the protospacers assigned by CRISPick within the chosen region. To minimize off-target effects, the selected protospacer sequences were screened against the *Z. tritici* IPO323 genome using BLASTn. sgRNAs were retained only if no more than 15 nucleotides outside the target gene matched the protospacer sequence, ensuring specificity [32].

In vitro transcription of sgRNAs was performed by generating forward primers with a T7 promoter, followed by two G nucleotides, the target-specific region, and a constant sequence. DNA templates for transcription were generated by annealing specific and constant oligonucleotides, followed by strand extension using T4 DNA polymerase. Annealing was carried out in a thermocycler with the following conditions: 95 °C for 5 min, gradual cooling from 95 °C to 85 °C at 2 °C/sec, and from 85 °C to 25 °C at 0.1 °C/sec. The annealed oligos were incubated with the T4 DNA mix at 12 °C for 20 minutes. The products were purified using NucleoSpin Gel and PCR clean-up (Macherey & Nagel GmbH & Co. KG, Düren, Germany).

The concentration of the DNA template size was measured with a Nanodrop spectrophotometer and the size was verified on a 3% agarose gel. sgRNAs were transcribed *in vitro* using the HiScribe T7 High Yield RNA Synthesis Kit (New England Biolabs, Ipswich, MA, USA, E2040L), following the manufacturer’s instructions with few modifications. After transcription, sgRNAs were purified using the RNA Clean & Concentrator-25 kit (Zymo Research, Irvine, CA, USA, R1018) according to the manufacturer’s instructions. The sgRNA concentration was measured using a Nanodrop, and purified sgRNAs were stored at -80° C.

### RNP formation and *in vitro* cleavage

Five sets of primers were designed to amplify the target DNA regions of the selected candidate effector genes Mycgr3106502, Mycgr3109710, Mycgr3107904, Mycgr3103393, and Mycgr394190 (Table 1). The resulting PCR amplicons were observed on a 1% agarose gel and purified using the NucleoSpin Gel and PCR Clean-up Kit (Macherey & Nagel) for the subsequent testing of Cas9/sgRNA cleavage.

**Table 1.**
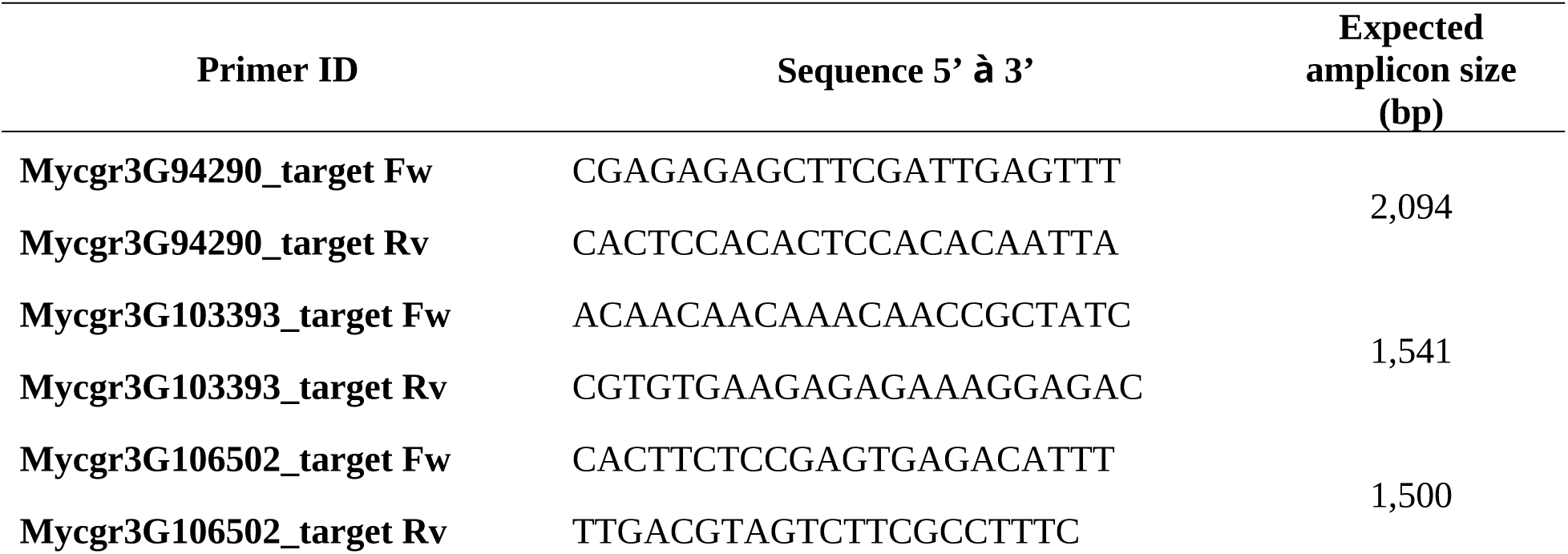
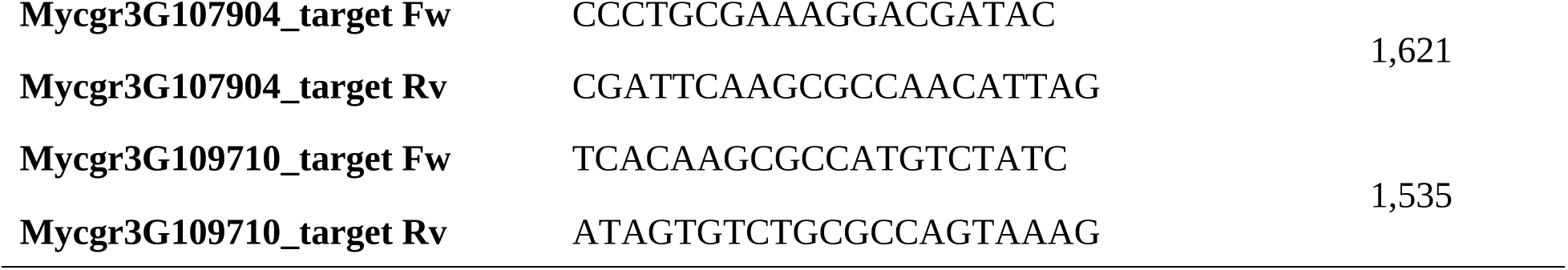
Primer sequences for amplification of the target DNA of five effector genes from the wheat pathogen *Zymoseptoria tritici*.

Ribonucleoprotein (RNP) complexes were assembled in vitro using *Streptococus pyogenes* Cas9 Nuclease (20 mM) (New England Biolabs, Ipswich, MA, USA, M0386T). The enzyme was diluted to 15 mM by mixing 7.5 µL of Cas9 with 2.5 µL of Diluent B (New England Biolabs, B8533S, Ipswich, MA, USA). Each RNP complex was assembled in a final volume of 10.5 µL containing 2 µL of DEPC-treated water, 2 µL of cleavage buffer r3.1 (New England Biolabs, Ipswich, MA, USA, B6003S), 1 µL of Cas9 enzyme (15 mM), and 1 µL of sgRNA (2,000 ng/µL, or 2 µg total). The reaction mixture was incubated at room temperature for 15 minutes to allow RNP formation. Cleavage reactions were initiated by adding 4 µL of purified target DNA (100 ng/µL) to the RNP complex, followed by incubation at 37 °C for 1.5 hours. To terminate the reaction, 1.0 µL of sterile EDTA (pH 8.0, Invitrogen, Waltham, MA, USA, AM9260G) was added, followed by 1.0 µL of Proteinase K (10 mg/mL) (Thermo Scientific, Waltham, MA, USA, EO0491) to denature the Cas9 protein. The reaction was then incubated at 37 °C for 20 minutes. Cleavage efficiency was assessed on a 2% agarose gel. A control reaction was included for one of the candidate effector genes, omitting the sgRNA while keeping all other components, to confirm that Cas9 alone does not cleave the target DNA.

### Construction of the donor DNA

The donor DNA template, containing the hygromycin B (HygB) resistance cassette *Pcpc1-hph-T-trpC* (*hph* under the control of the *Neurospora. crassa pCPC1* and *Aspergillus nidulans tTrpC*) promotors [48], was amplified from plasmid pTel_HygR [32]. Five sets of primers were designed for amplification, each incorporating 60-bp regions matching the immediate upstream and downstream sequences of the target effector genes. Primers included the universal binding sequences TL39 (5′-TGCTGGCCTTTTGCTCACATGCATG**-**3′) and TL40 (5′-ATCGCCGGAAAGGACCCGCAAATG**-**3′) in pTel_HygR (Supplementary Table 1). For the reverse primers, the 60-bp matching regions were used in their reverse-complement orientation (See Supplementary Table 2 for complete sequences of primers).

The HygB cassette was amplified by PCR using a reaction mixture containing 0.5 µL of each forward and reverse primer, 0.5 µL of 10 mM dNTPs, 0.5 µL of pTel_Hyg plasmid, 0.25 µL of Phusion Plus Polymerase, and 17.75 µL of DEPC-treated H₂O, for a total volume of 25 µL. The PCR conditions were: initial denaturation at 98 °C for 30 sec, followed by 35 cycles of 98 °C for 10 sec, 60 °C for 10 sec, and 72 °C for 50 sec, with a final extension at 72 °C for 5 min and an indefinite hold at 4 °C. Amplification products were verified by electrophoresis on a 1% agarose gel stained with SybrSafe (Invitrogen, S33102). The expected 2,500-bp DNA fragments for the five donor DNA templates were purified using the NucleoSpin Gel and PCR Clean-up Kit (Macherey & Nagel). Multiple rounds of purification, utilizing at least two columns, were performed to concentrate the donor DNA, yielding a final amount of 5000 ng (5 µg) per transformation. The purified donor DNA was then diluted in 60 µL of Tris-EDTA buffer (Sigma-Aldrich, 93283) with 40 mM CaCl₂.

### Transformation of *Z. tritici*

Transformation was performed following a published protocol for *Botrytis* cinerea [49], optimized by Steentjes [33] for *P. fijiensis* and adapted for *Z. tritici*. A volume of 0.7 mL of protoplast solution was used per experiment, containing approximately 26.2 × 10⁶ protoplasts. To standardize the number used for each transformation, the total count was divided by seven, yielding 3.75 × 10⁶ protoplasts per 100 µL, which was used for each of the seven constructs, including five candidate genes, positive and negative controls. Protoplasts (100 µL or 3.75 × 10⁶ protoplasts) were transformed with one RNP complex per gene (15 uL, 2,000 ng of the selected sgRNA) and 60 uL of purified donor template DNA (5,000 ng) containing the HygB cassette.

The DNA and RNP mixture was added to the protoplast suspension. The samples were incubated on ice for 10 minutes. An equal volume of 175 µL of PEG 3350 (Sigma-Aldrich, 88276) solution was then added to each sample using a cutoff blue pipette tip. The mixture was gently pipetted up and down 20–30 times to ensure proper mixing and incubated at room temperature for 20 minutes. To facilitate recovery of the transformed protoplasts, 680 µL of TMSC buffer (18.2 g of D-Sorbitol (1 M), 209 mg of MOPS, 588 mg of CaCl_2_ in 100 mL of milliQ H_2_O, pH 6.3) at room temperature was added, followed by centrifugation at 1,500 × g for 3 minutes in a swing-out rotor and an additional 3 minutes at 3,000 rpm in a standard microcentrifuge. The supernatant was carefully removed, and the pellet was resuspended in 200 µL of TMSC buffer.

The transformed protoplast suspension was transferred into 25 mL of liquid RM agar (42 °C), mixed thoroughly, and poured into two, 90-mm petri dishes (12.5 mL per plate) without a selection agent. Plates were incubated overnight (18 hours) at 18 °C. The following day, a 12.5-mL overlay of RM agar containing Hygromycin B (200 µg/mL) was added to each plate (except the positive controls) so that each plate with a total volume of 25 mL of RM agar had a final concentration of 100 µg/mL HygB. Subsequently, plates were incubated at 18 °C to allow for selection and inspected visually after 10, 12 and 20 days. The positive control consisted of protoplasts grown on antibiotic-free medium without Cas9, sgRNA, or donor template to test the regeneration of the protoplasts. The negative control consisted of protoplasts grown on medium containing the same HygB concentration as the transformed protoplasts but without Cas9, sgRNA, or donor template, in which no colonies were expected to grow.

### PCR analysis of putative *Z. tritici* effector knockouts

Genomic DNA was extracted from the *Z. tritici* wild-type strain IPO323 and mutant colonies (*Z. tritici* ΔMycgr3109710, ΔMycgr394290, ΔMycgr3107904_1, and ΔMycgr3107904_2) using the Master Pure Yeast DNA Purification Kit (Biosearch Technologies, Middleton, WI, USA; MPY80200). DNA concentration and quality were assessed with a NanoDrop spectrophotometer. To confirm insertion of the donor DNA construct and disruption of the target genes, PCR amplification was performed using primers located approximately 500 bp upstream and downstream of each target gene. For the ΔMycgr3107904_1 and ΔMycgr3107904_2 mutants, an additional PCR reaction for outward amplification was conducted using two internal primers for the HygB cassette (GTATGAGTCACAGCACCGATAC and CGGGTTTACCTCTTCCAGATAC) in combination with the forward or reverse primers for Mycgr3107904 to determine the insertion orientation of the HygB cassette. PCR products were analyzed by electrophoresis on a 1% agarose gel stained with SybrSafe (Invitrogen,; S33102), purified using the NucleoSpin Gel and PCR Clean-up Kit (Macherey & Nagel), and sequenced by Sanger sequencing (Supplementary data).

### Assessment of colony morphology in the putative knockout strains

The growth phenotypes of *Z. tritici* mutants were assessed on PDA, YSA, yeast-maltose agar (YMA), RM, and water agar (WA) plates. A 10-µL drop of conidial suspension (∼2 × 10⁷ conidia/mL) was placed at the center of each plate. The plates were incubated at 18 °C and monitored visually at 7 and 14 days postinoculation.

### Phenotypic assessment of virulence caused by knockout strains

For virulence assays, conidia from the WT strain IPO323 and the two Mycgr3107904 mutant strains (ΔMycgr3107904_1 and ΔMycgr3107904_2) were collected by centrifugation from 4-day-old liquid YSB cultures. The conidia were resuspended in sterilized milliQ water, adjusted to a concentration of 1 × 10⁷ conidia/mL, and supplemented with three drops of Tween20.

Fourteen-day-old seedlings of the susceptible wheat cultivar Taichung29 were inoculated with either the *Z. tritici* IPO323 wild-type strain, the Δ*Mycgr3107904_1* or Δ*Mycgr3107904_2* mutant strains using the spray-inoculation method [50]. For each strain, 14 to 16 wheat plants were inoculated, with the second leaf labeled for scoring across multiple time points. An uninoculated control group of 14 plants was included. Disease progression was assessed by measuring the symptomatic leaf area at 7, 10, 12, 14, and 19 days post inoculation (DPI). At each time point, three plants per treatment were evaluated, and the experiment was conducted twice to ensure reproducibility. Chlorotic and necrotic lesions were quantified using ImageJ, and the percentage of affected leaf area was calculated. A Tukey’s test was performed to determine statistically significant differences in disease severity between the WT strain and the Mycgr3107904 knockout strains.

## Results

### Generation of protoplasts from *Z. tritici* blastospores

Because efficient protoplast generation is essential for CRISPR/Cas9-based genome editing in *Z. tritici*, we first sought to identify culture conditions that promote robust blastospore production. We first assessed the growth of *Z. tritici* IPO323 on three liquid culture media at 18 °C after one to two days of growth from a four-day-old colony growing on PDA or YSA. On MSB, IPO323 primarily developed hyphae, with limited blastospore formation and terminal buds observed after two days (Figure 1). In contrast, growth on PDB resulted in a higher number of budding points on germinated pycnidiospores, which are crucial for blastospore production [51]. PDB cultures contained a higher number of free blastospores (Figure 1). Since newly formed cells have less reinforced cell wall, they are presumably more sensitive to enzymatic digestion, and therefore they may be more suitable as a primary source for protoplast formation.

**Figure 1.**
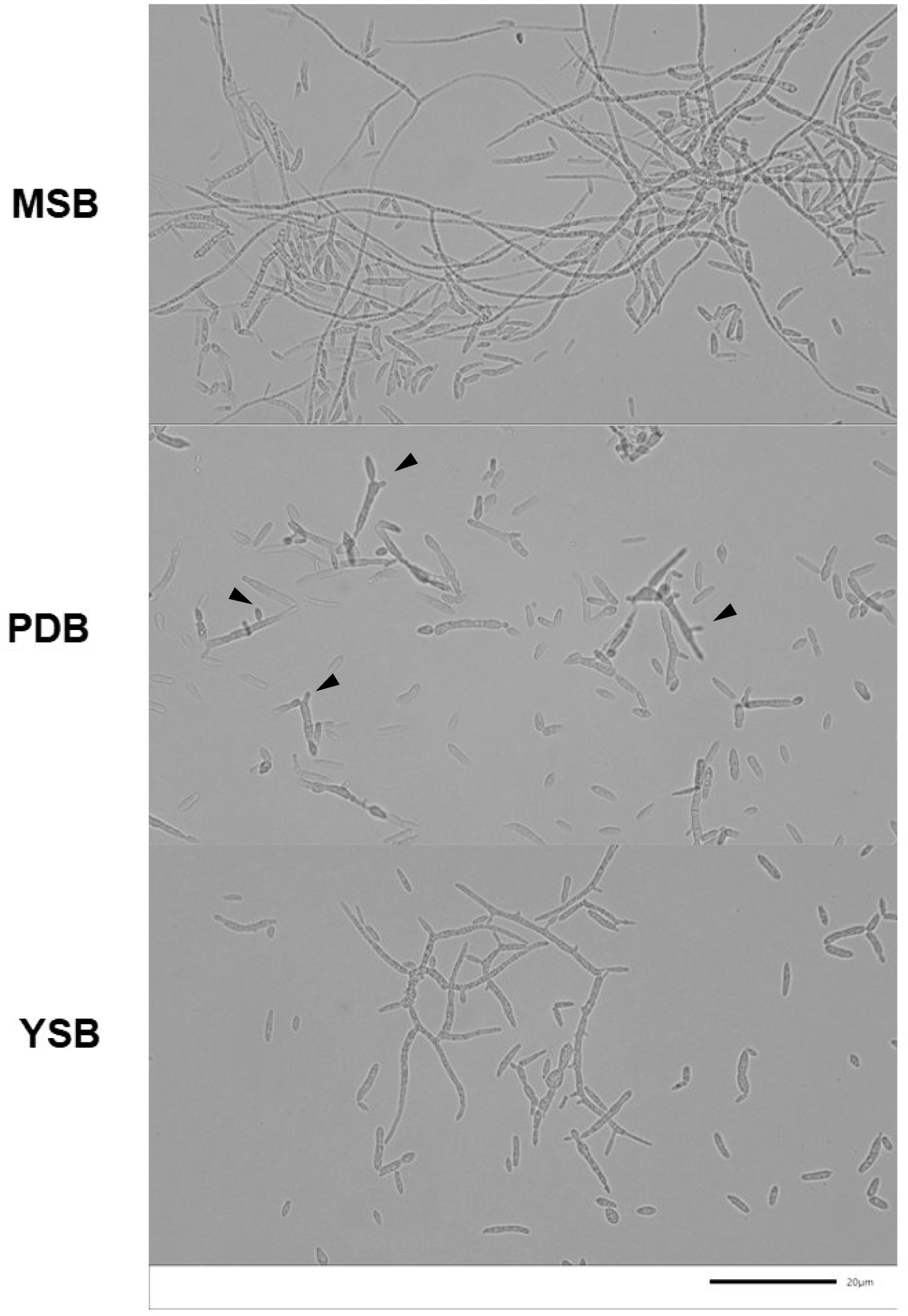
Microscopic observation of *Zymoseptoria tritici* growth in three liquid media after one day of incubation. MSB: Maltose sucrose broth; PDB: Potato dextrose broth; YSB: Yeast sucrose broth.

Growth in YSB showed some hyphal branching, though less extensive than in MSB, and a higher frequency of terminal buds compared to MSB, but lower than in PDB (Figure 1).

Ultimately, PDB yielded approximately 5.7 x 10⁷ conidia/mL, with a higher proportion of free blastospores, making it the most suitable medium for growth of *Z. tritici* to obtain fungal biomass for protoplast formation.

A 24-hour old liquid culture of *Z. tritici* grown in PDB at 18 °C was used as the starting material for enzymatic digestion. After 40 minutes at 28 °C, the first protoplasts appeared as rounded structures (Figure 2), alongside intact blastospores and cell debris that were likely derived from germinated or enlarged pycnidiospores and blastospores. After 70 minutes, protoplast numbers had increased (Figure 2), and their concentration was determined using a hemocytometer, yielding 37.5 × 10⁶ protoplasts/mL. Extending digestion beyond 70 minutes resulted in protoplast lysis and increased debris, indicating 70 minutes was the optimal digestion time (Figure 2).

**Figure 2.**
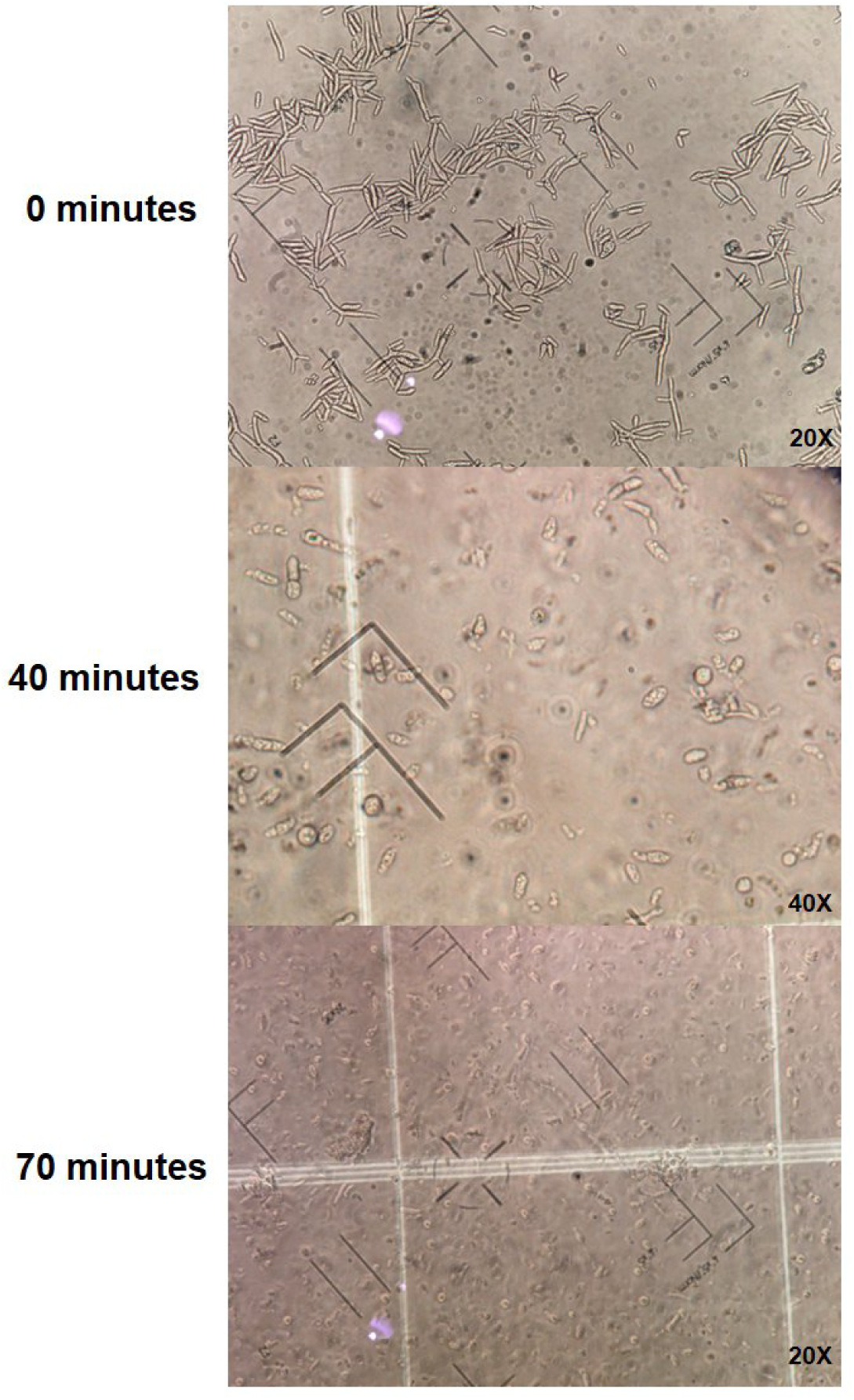
Protoplast formation of *Zymoseptoria tritici* at three time points during enzymatic digestion: initiation or 0 minutes; 40 minutes; and 70 minutes after digestion began.

### *In vitro* cleavage assay and donor DNA template

To delete five candidate effector genes in IPO323, we designed and selected two sgRNAs per gene with the highest on-target efficiency scores as predicted by CRISPick, and with no off-target matches identified by *in silico* BLASTn analysis (Table 2). All selected sgRNAs have fewer than 15 nucleotide matches with other genomic regions in the *Z. tritici* IPO323 reference genome (GenBank accession number: GCF_000219625.1).

**Table 2.**
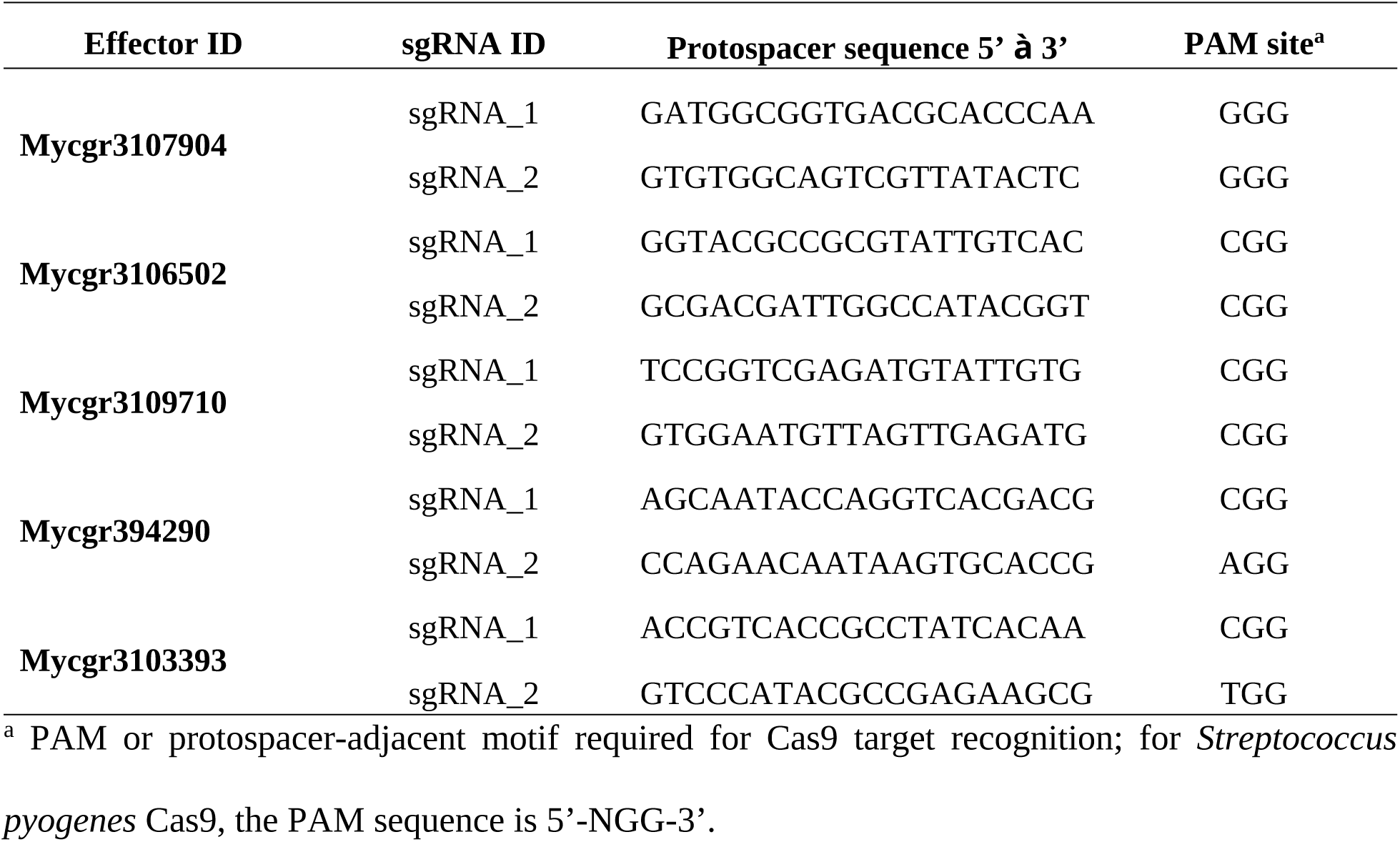
sgRNAs targeting putative *Zymoseptoria tritici* effector genes for transformation.

Synthesis of sgRNA1 for *Mycgr3109710* and sgRNA2 for *Mycgr349290* was repeated three times without achieving the required 2,000-μg concentration for Cas9/sgRNA complex formation and cleavage testing. As a result, only one sgRNA was used for cleavage assays on these effector genes.

Functionality of *in vitro-*assembled Cas9-sgRNA complexes was tested by cleavage of target DNA *in vitro*. First, the target DNA regions of the five candidate genes were amplified using primers that annealed within the flanking regions, covering 500 bp upstream and downstream of the target sites (Table 1). PCR amplification confirmed the expected product sizes for all target genes (Table 1, Figure 3).

**Figure 3.**
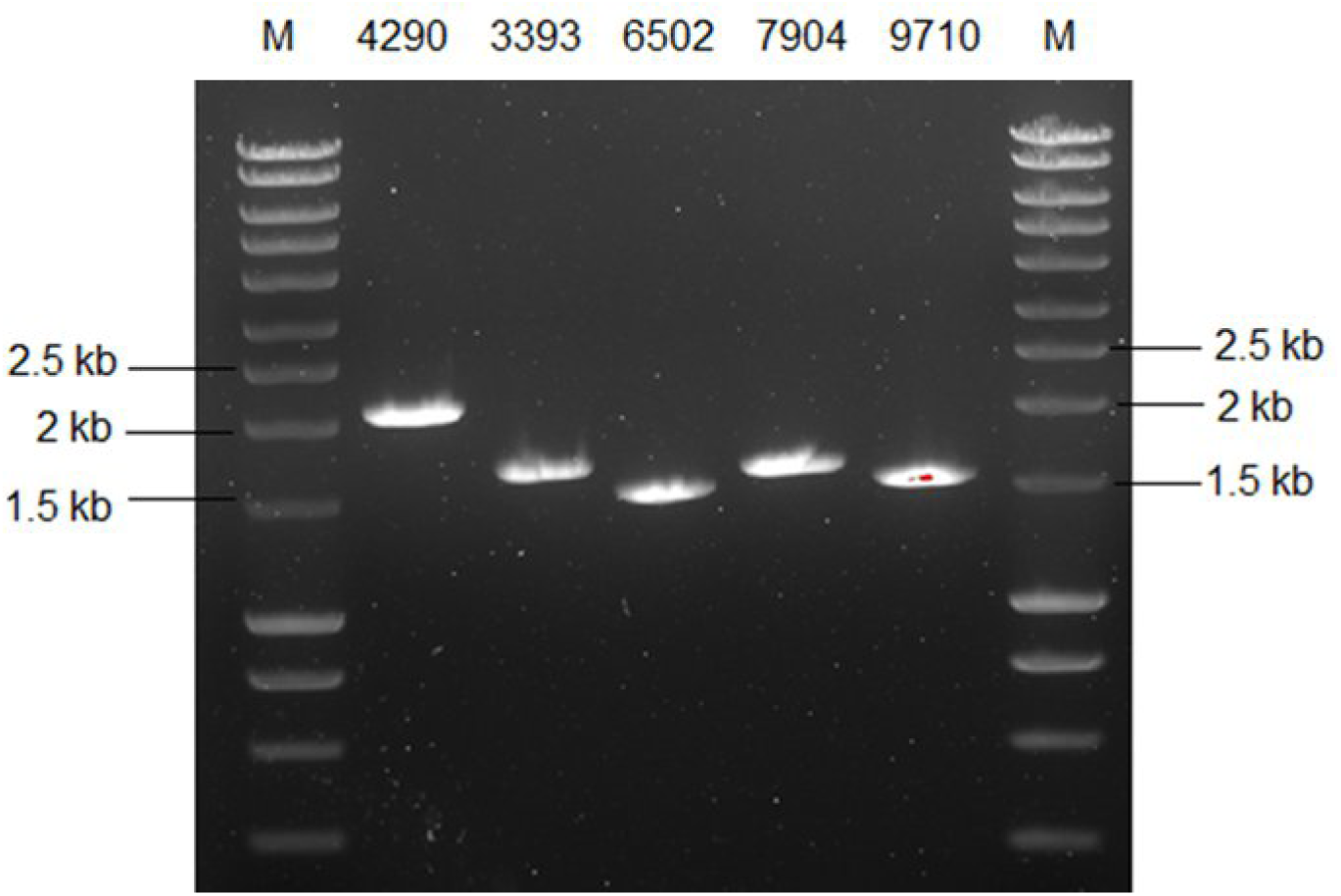
PCR amplification of target DNA for five candidate effector genes from the wheat pathogen *Zymoseptoria tritici*. M: 10-kb SmartLadder. Expected amplicon sizes for *Mycgr394290*, *Mycgr3103393*, *Mycgr3106502*, *Mycgr3107904*, and *Mycgr3109710* was 2,094, 1,541, 1,500, 1,621, and 1,535 bp, respectively.

PCR-amplified fragments of the coding regions of the five candidate effector genes were mixed with their respective RNP complexes consisting of the pre-mixed sgRNA and Cas9 protein to assess cleavage efficiency. Cas9-mediated cleavage was evaluated by agarose gel electrophoresis (Figure 4). For *Mycgr3107904*, the Cas9-sgRNA2 complex successfully cleaved the entire template within 90 minutes, producing fragments of the expected size (1,064 and 567 bp). Similarly, for *Mycgr3103393*, both Cas9-sgRNA complexes efficiently cleaved nearly all the template within the same timeframe. For *Mycgr3109710* and *Mycgr394290*, the single Cas9-sgRNA complex tested for each gene also achieved near-complete cleavage within 90 minutes.

**Figure 4.**
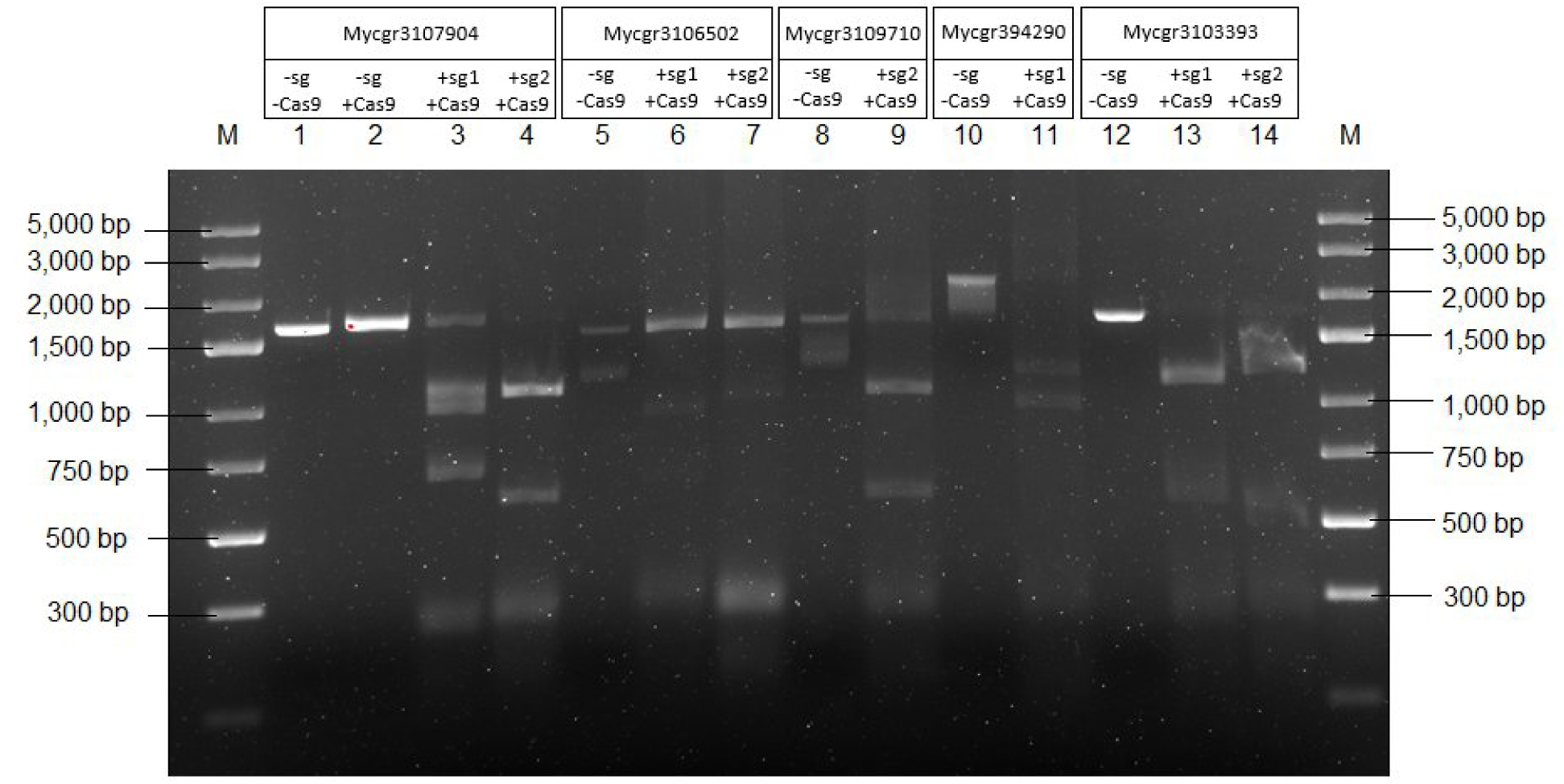
Cleavage of sgRNAs on amplified regions of five candidate effector genes from the wheat pathogen *Zymoseptoria tritici*. M: 5-kb Ladder; 1 – 4, Target DNA of Mycgr3107904: 1 = no sgRNA, no Cas9; 2 = no sgRNA, Cas9; 3 = sgRNA1 + Cas9; 4 = sgRNA2 + Cas9; 5 – 7, Target DNA of Mycgr3106502: 5 = no sgRNA, no Cas9; 6 = sgRNA1 + Cas9; 7 = sgRNA2 + Cas9; 8 – 9, Target DNA of Mycgr3109710: 8 = no sgRNA, no Cas9, 9 = sgRNA2 + Cas9; 10 – 11, Target DNA of Mycgr394290: 10 = no sgRNA, no Cas9; 11 = sgRNA1 + Cas9; 12 – 14, Target DNA of Mycgr3103393: 12 = no sgRNA, no Cas9; 13 = sgRNA1 + Cas9; 14 = sgRNA2 + Cas9.

However, for *Mycgr3106502*, neither of the two Cas9-sgRNA complexes fully cleaved the template. Despite the lower efficiency, both produced cleaved fragments of the expected size, leading to the selection of one sgRNA for gene knockout attempts.

The RNP complexes that were eventually used for transformation of *Z. tritici* protoplasts were assembled using one sgRNA per gene, selected based on Cas9/sgRNA cleavage efficiency. The sgRNAs used were as follows: sgRNA2 for *Mycgr3107904*; sgRNA1 for *Mycgr3106502*; sgRNA2 for *Mycgr3109710;* sgRNA1 for *Mycgr394290;* and sgRNA1 for *Mycgr3103393*.

The donor DNA template was amplified, yielding fragments of the expected size (∼2,500 bp) for all five targeted effector genes (Figure 5). These 2,500 bp include the 2,380-bp HygB cassette, amplified from the TL39 to the TL40 universal binding sites, along with an additional 120 bp of homologous regions flanking the target genes (60 bp upstream and 60 bp downstream).

**Figure 5.**
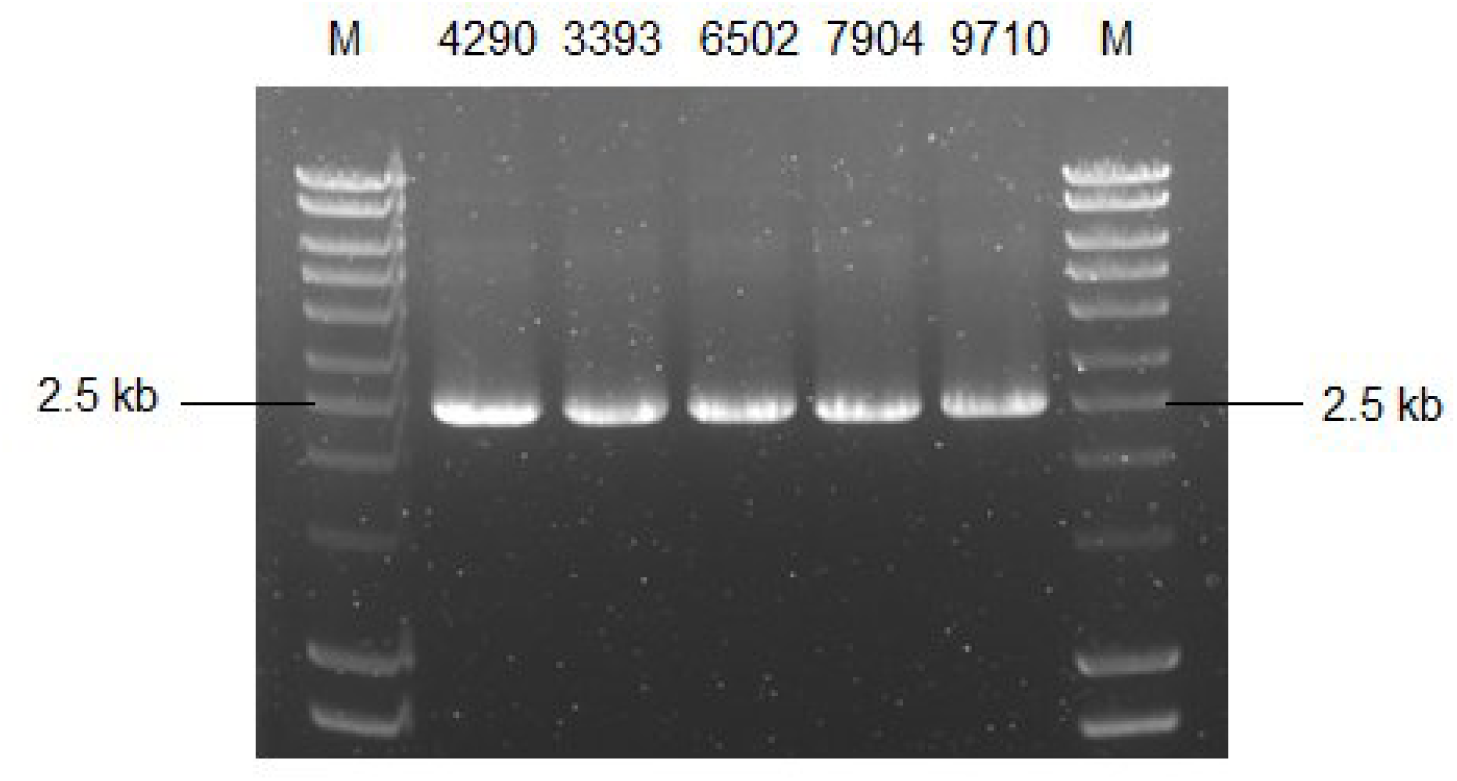
PCR amplification of the donor DNA templates containing the amplified HygB cassette from pTel_HygB and 60 bp with complementarity to upstream and downstream regions of the target gene sequences for the five selected putative effector genes Mycgr394290, Mycgr3103393, Mycgr3106502, Mycgr3107904, and Mycgr3109710 of the wheat pathogen *Zymoseptoria tritici*.

### Generation of deletion mutants for candidate effector genes in *Z. tritici* via Cas9-mediated transformation

After the transformation, protoplasts were regenerated in selective medium for 20 days, and two putative transformant colonies were obtained for the effector gene *Mycgr3107904*, while a single putative transformant was obtained for each of *Mycgr394290* and *Mycgr3109710*. No colonies were observed for protoplasts growing on selective medium (100 µg/mL hygromycin) for *Mycgr3106502* or *Mycgr3103393*. No colonies grew on the negative control (medium with 100 ug/mL of hygromycin) plates, and colonies were observed on the positive control plates (medium without hygromycin), indicating the ability of the protoplasts to regenerate (Supplementary figure 1).

### Phenotyping of *Z. tritici* mutants

Fresh liquid cultures of the transformants were used to prepare conidial suspensions at a concentration of approximately 2 × 10⁷ conidia/mL. A 10-µL drop of each suspension was inoculated at the center of a plate, and colony morphology was evaluated at 7 and 14 days post inoculation. By day 7, the *Z. tritici* Δ*Mycgr3107904_1* mutant exhibited a distinct colony morphology compared to the wild type on YSA, PDA, and YMA (Figure 6). The mutant displayed a faster growth rate, surpassing the colony size of the wild type and other transformants. Notably, Δ*Mycgr3107904_1* showed no signs of pigmentation or hyphal growth, whereas the wild-type strain exhibited hyphal development and pigmentation, particularly along the colony edges, with the most pronounced hyphal growth observed on PDA and YMA (Figure 6).

**Figure 6.**
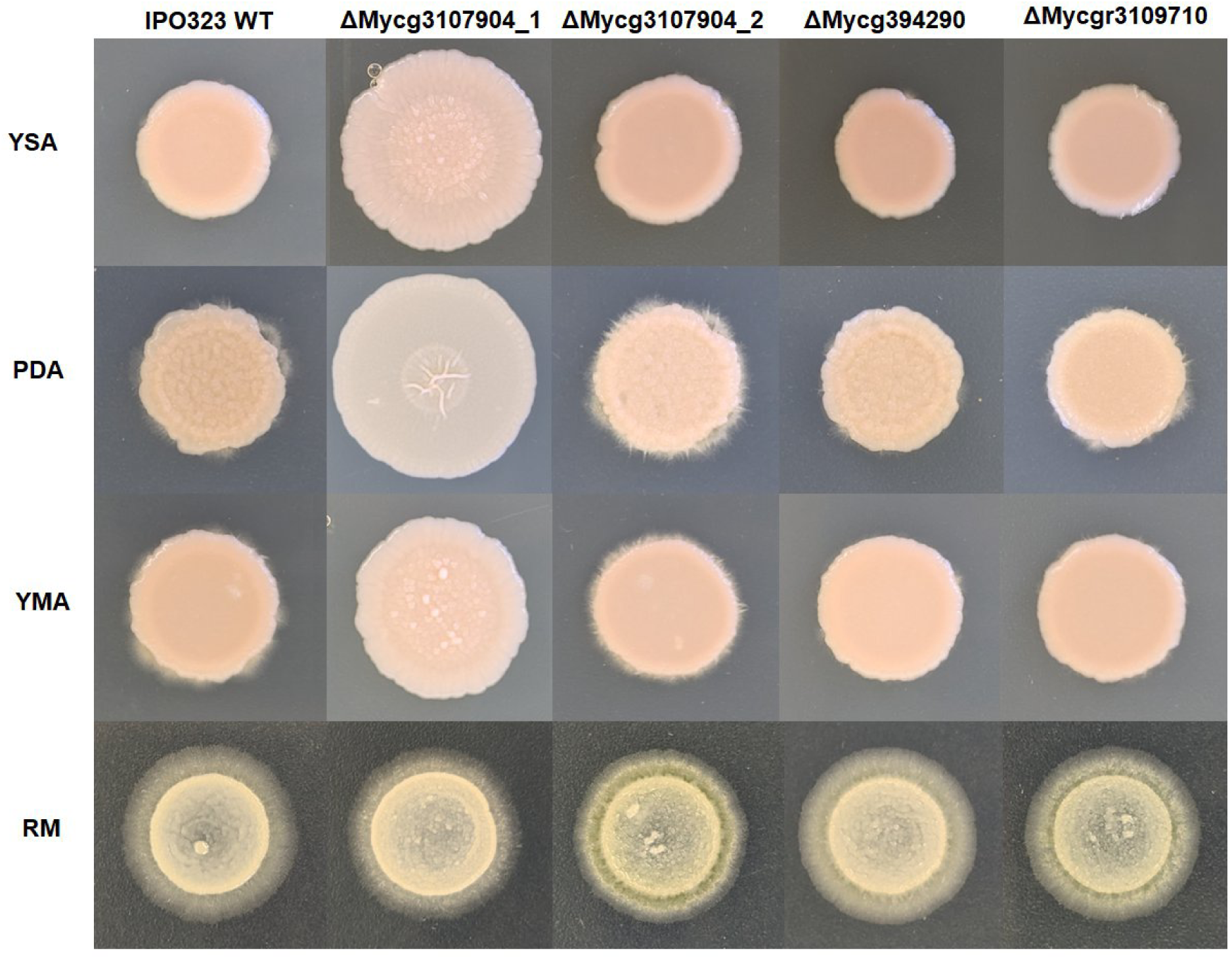
Growth on four types of medium at 7 days after inoculation for the *Zymoseptoria tritici* WT IPO323 strain and the hygromycin-resistant strains ΔMycgr3107904_1, ΔMycgr3107904_2, ΔMycgr394290, and ΔMycgr3109710. YSA: Yeast sucrose agar; PDA: Potato dextrose agar; YMA: Yeast maltose agar; RM: Regeneration medium.

At 14 days post inoculation, the Δ*Mycgr3107904_1* putative mutant showed no signs of hyphal growth on YSA, PDA, or YMA and exhibited significantly reduced hyphal development on water agar (WA) compared to the wild-type strain (Figure 7). WA is commonly used to induce hyphal growth in *Z. tritici* [52]. On PDA, Δ*Mycgr3107904_1* displayed a noticeable change in pigmentation, with darkening originating from the center, suggesting the onset of melanization (Figure 7). This mutant also maintained a faster growth rate, forming a larger colony diameter than the wild type and other transformants.

**Figure 7.**
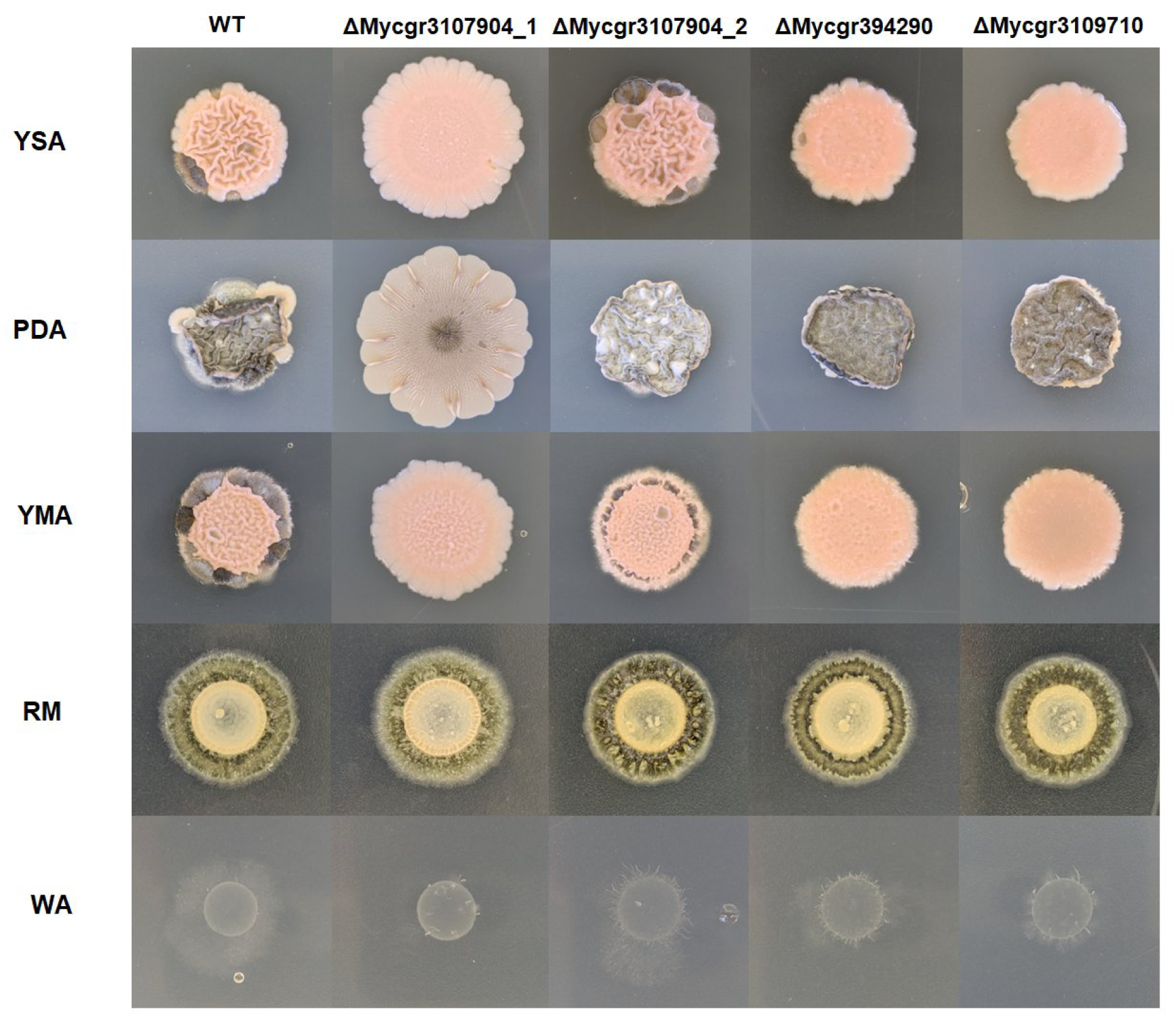
Growth on four types of medium at 14 days after inoculation for the *Zymoseptoria tritici* WT IPO323 strain, and the hygromycin-resistant strains ΔMycgr3107904_1, ΔMycgr3107904_2, ΔMycgr394290, and ΔMycgr3109710. YSA: Yeast sucrose agar; PDA: Potato dextrose agar; YMA: Yeast maltose agar; RM: Regeneration medium; WA: Water agar.

In contrast, the colony morphology of Δ*Mycgr3107904_2* resembled more that of the wild type, although with differences in pigmentation on PDA and less hyphal development on YMA. Mutants Δ*Mycgr394290* and Δ*Mycgr3109710* looked similar to each other and did not exhibit melanization or hyphal growth in contrast to the wild-type strain at 14 days post inoculation on YSA and YMA, indicating a significant delay compared to the wild-type strain. Additionally, all four putative mutants showed markedly reduced hyphal development on WA, further distinguishing them from the wild type (Figure 7).

### Genotyping of *Z. tritici* mutants

The putative *Z. tritici* mutants obtained from selective media were screened initially by PCR using primers designed in the upstream and downstream flanking regions of the target genes. This approach allowed detection of *HygB* cassette insertion and potential differences in fragment sizes resulting from DSB repair. PCR amplification was performed using DNA from the wild-type strain as a control and from purified cultures of *HygB*-resistant *Z. tritici* mutants. For putative mutant Δ*Mycgr394290*, a ∼2,000-bp DNA fragment was obtained, which is shorter than the corresponding wild-type amplicon (Figure 8A). This suggests that nucleotide deletions may have occurred during DSB repair, while the *HygB* cassette was ectopically inserted, conferring hygromycin resistance.

**Figure 8.**
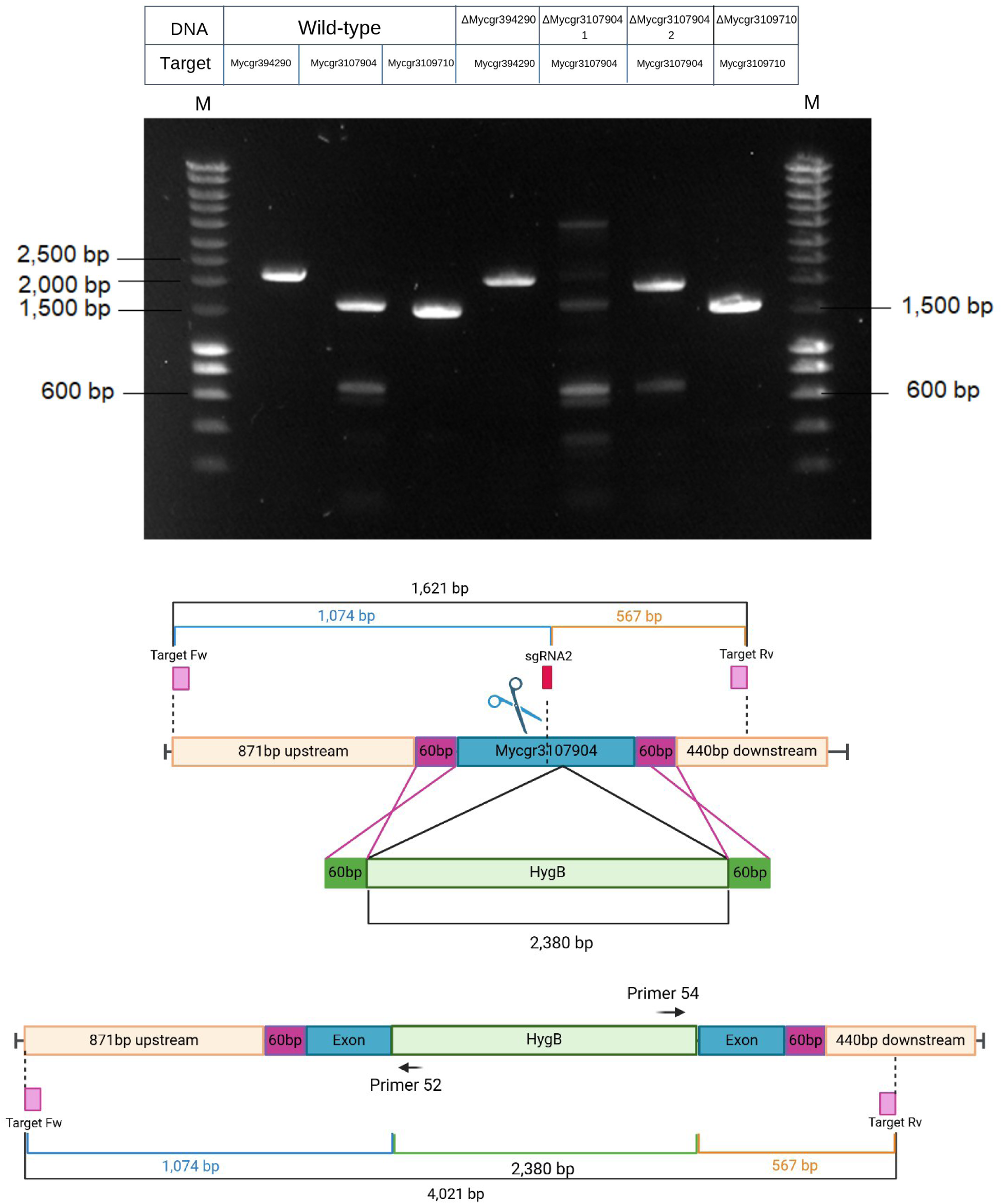
PCR amplifications of putative *Zymoseptoria tritici* CRISPR mutants and the wild-type strain using primers for the target DNA regions and a cartoon of the expected insertion for one mutant as an example. A) M = 10-kb marker ladder (Smartladder); 1 – 3, WT DNA. 1: Mycgr394290 (2,094 bp); 2: Mycgr3107904 (1,621 bp); 3: Mycgr3109710 (1,535 bp); 4: ΔMycgr394290 DNA, amplicon is ∼2,000 bp; 5: ΔMycgr3107904_1 DNA, two DNA fragments of ∼1,500 bp and ∼4,000 bp were obtained; 6: ΔMycgr3107904_2 DNA, amplicon is ∼1,900 bp; 7: ΔMycgr3109710 DNA, amplicon is ∼1,550 bp. B) Schematic representation of the *Mycgr3107904* gene disruption by Cas9-mediated transformation coupled with microhomology-directed repair (MDR). The cleavage site of the *in vitro*-assembled Cas9/sgRNA2, the 60-bp microhomology regions for MDR, the location of the forward and reverse primers for amplification of the target gene, and the expected sizes of each region are represented for the genomic locus of the wild-type IPO323 and the Δ ΔMycgr3107904_1 transformant strain. C) Schematic representation of the expected *Mycgr3107904* gene disruption following insertion of the HygB gene.

For putative mutant Δ*Mycgr3107904_1*, two distinct PCR fragments were observed: one ∼4,000-bp fragment (Figure 8A), consistent with *HygB* cassette insertion at the predicted cleavage site of sgRNA2 (Figure 8B), and another fragment similar in size to the wild-type amplicon. This suggests the presence of a mixed population, potentially including an ectopic mutant recovered from the selective medium. Further colony purification is necessary to confirm this hypothesis. For putative mutant Δ*Mycgr3107904_2*, a single ∼1,900-bp fragment was detected, larger than the wild-type amplicon. This indicates that, although the *HygB* cassette likely was inserted ectopically, additional sequences may have been incorporated during the repair process, potentially disrupting the target gene. Finally, for putative mutant Δ*Mycgr3109710*, the PCR product was slightly larger than the wild-type fragment (Figure 8A), suggesting nucleotide alterations along with ectopic *HygB* cassette insertion.

Further PCR screening was performed to confirm the correct integration of the *HygB* cassette into the *Mycgr3107904* locus via MDR. To determine the orientation of the insertion, PCR amplification was conducted using two internal primers that amplify from within the *HygB* cassette into the flanking regions in combination with the gene-specific primers for *Mycgr3107904*. If the cassette is integrated in the 5’ to 3’ orientation, we expect amplification with Mycgr3G107904_target Fw + primer 52 and primer 54 + Mycgr3G107904_target Rv (Figure 9A). If inserted in the 3’ to 5’ reverse orientation, amplification is expected with Mycgr3G107904_target Fw + primer 54 and Mycgr3G107904_target Rv + primer 52 (Figure 9B). However, PCR using DNA from mutant ΔMycgr3107904_1 produced fragments from both combinations with primer 52, a fragment of ∼1,500 bp (Figure 9C, lane 1) using Mycgr3G107904_target Fw + primer 52, and a fragment of ∼1,000 bp (Figure 9C, lane 4) using primer 52 + Mycgr3G107904_target Rv, suggesting a different insertion pattern.

**Figure 9.**
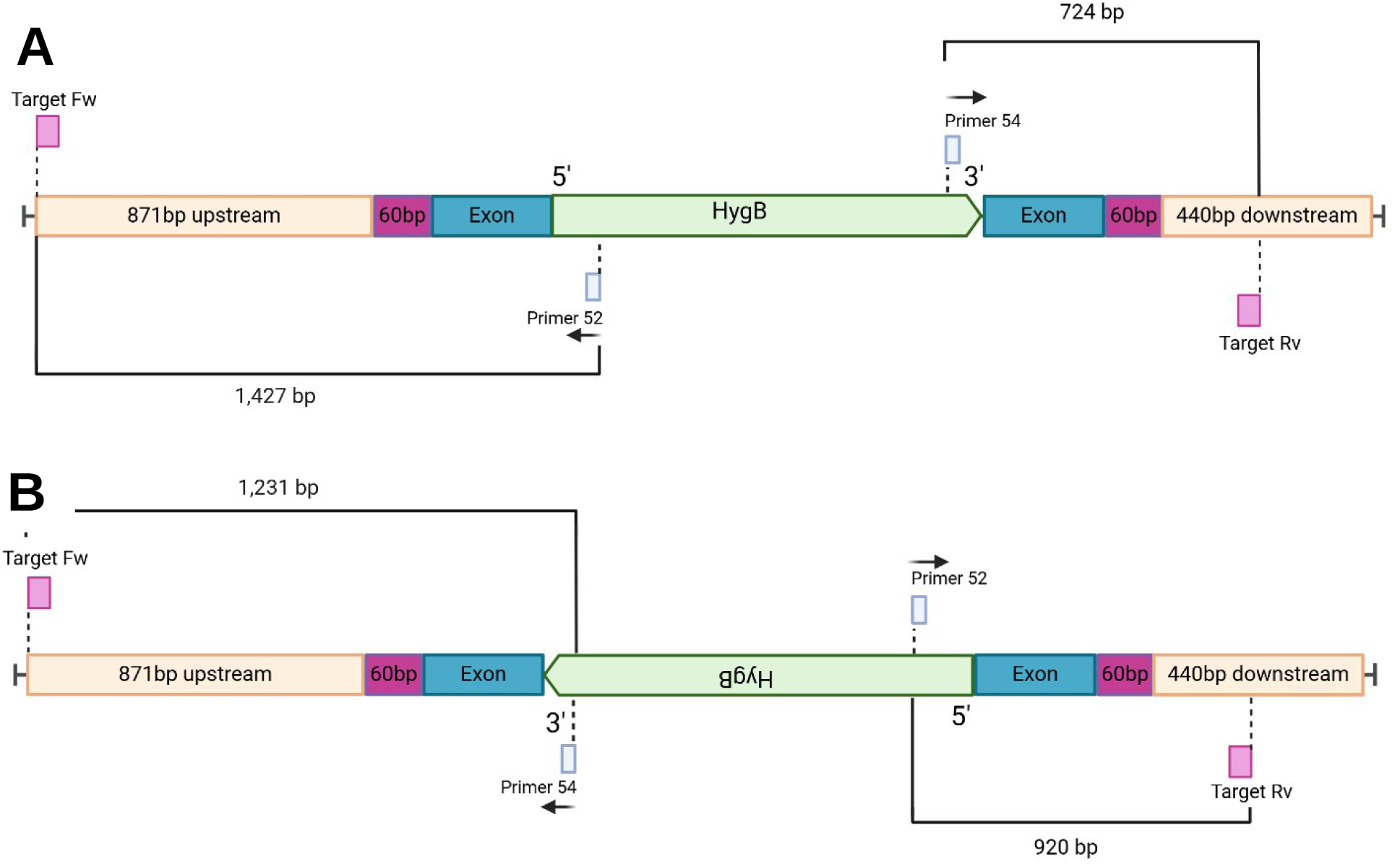
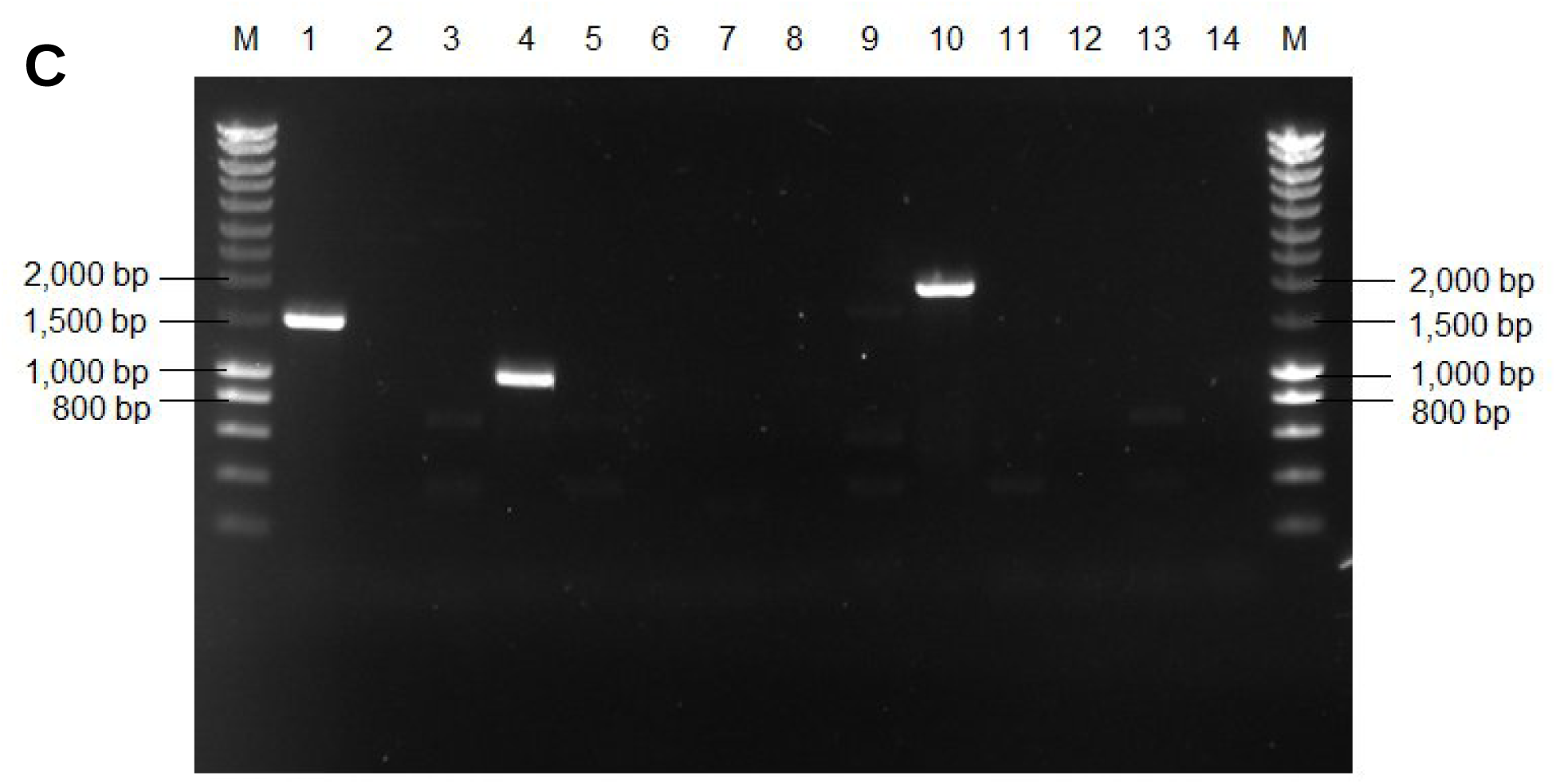
Schematic representation of two possible orientations of the HygB gene in *Z. tritici* mutants of the *Mycgr3107904* gene and PCR results with specific primer pairs. A) Insertion of the *HygB* cassette in the 5’ to 3’ orientation, or B) Insertion of the *HygB* cassette in the 3’ to 5’ orientation. C) PCR amplifications using combinations of the primers to amplify the target DNA regions and the internal primers for the HygB cassette. M = 10-kb SmartLadder; 1 – 4, 9: ΔMycgr3107904_1 DNA. 5 – 8, 10: ΔMycgr3107904_2 DNA. 11 – 14, WT DNA. Primers used for lanes 1, 5, 11: Mycgr3G107904_target Fw + primer 52; 2, 6, 12: primer 54 + Mycgr3G107904_target Rv; 3, 7, 13: Mycgr3G107904_target Fw **+** primer 54; 4, 8, 14: primer 52 + Mycgr3G107904_target Rv; 9 and 10: Mycgr3G107904_target Fw **+** Mycgr3G107904_target Rv.

For mutant ΔMycgr3107904_2, no bands were detected with any of the tested primer combinations using the internal primers for the *HygB* cassette (Figure 9C, lanes 5–8). However, amplification using the gene-specific primers for Mycgr3107904 produced a ∼1,900-bp fragment (Figure 9C lane 10), the same size observed in the initial PCR and larger than the wild-type amplicon. This suggests that the *HygB* cassette was not integrated, but nucleotide insertions likely occurred during DNA repair, disrupting the target gene. Sanger sequencing of the fragment in lane 10 (Figure 9C) (Supplementary data) confirmed this hypothesis, revealing two sequences: 1,028 bp and 287 bp, that cover regions of the target gene *Mycgr3107904* amplified from Mycgr3G107904_target Fw until the cleavage site of sgRNA2 and from Mycgr3G107904_target Rv, respectively. Additionally, a ∼298-bp sequence was sandwiched between them, but it did not overlap in the sequencing results from either gene-specific primer. This suggests that an even longer DNA fragment was inserted at the sgRNA2 cleavage site during DNA repair, disrupting *Mycgr3107904*.

To confirm specificity, we tested the same primer combinations with wild-type DNA (Figure 9C, lanes 11–14) and obtained no amplification, verifying that the fragments detected in mutant ΔMycgr3107904_1 were not false positives.

To further confirm the PCR amplifications for mutant ΔMycgr3107904_1, we performed an additional screening using two replicates of each primer combination and DNA from two independent extractions from the same mutant. The amplified DNA fragments were purified and sequenced using Sanger sequencing with primers Mycgr3G107904_target Fw, Mycgr3G107904_target Rv, primer 52, and primer 54. Consistent with the initial screening, we obtained amplicons of ∼1,500 bp (from Mycgr3G107904_target Fw + Primer 52) and ∼1,000 bp (from primer 52 + Mycgr3G107904_target Rv) (Figure 10A). Notably, two additional faint bands of ∼3,600 bp and ∼3,100 bp also were detected (Figure 10A, lanes 2 and 3, lanes 6 and 7). These results suggest that the *HygB* cassette was inserted multiple times in both orientations. A schematic representation of this proposed insertion pattern, explaining the observed fragment sizes, is shown in Figure 10B.

**Figure 10.**
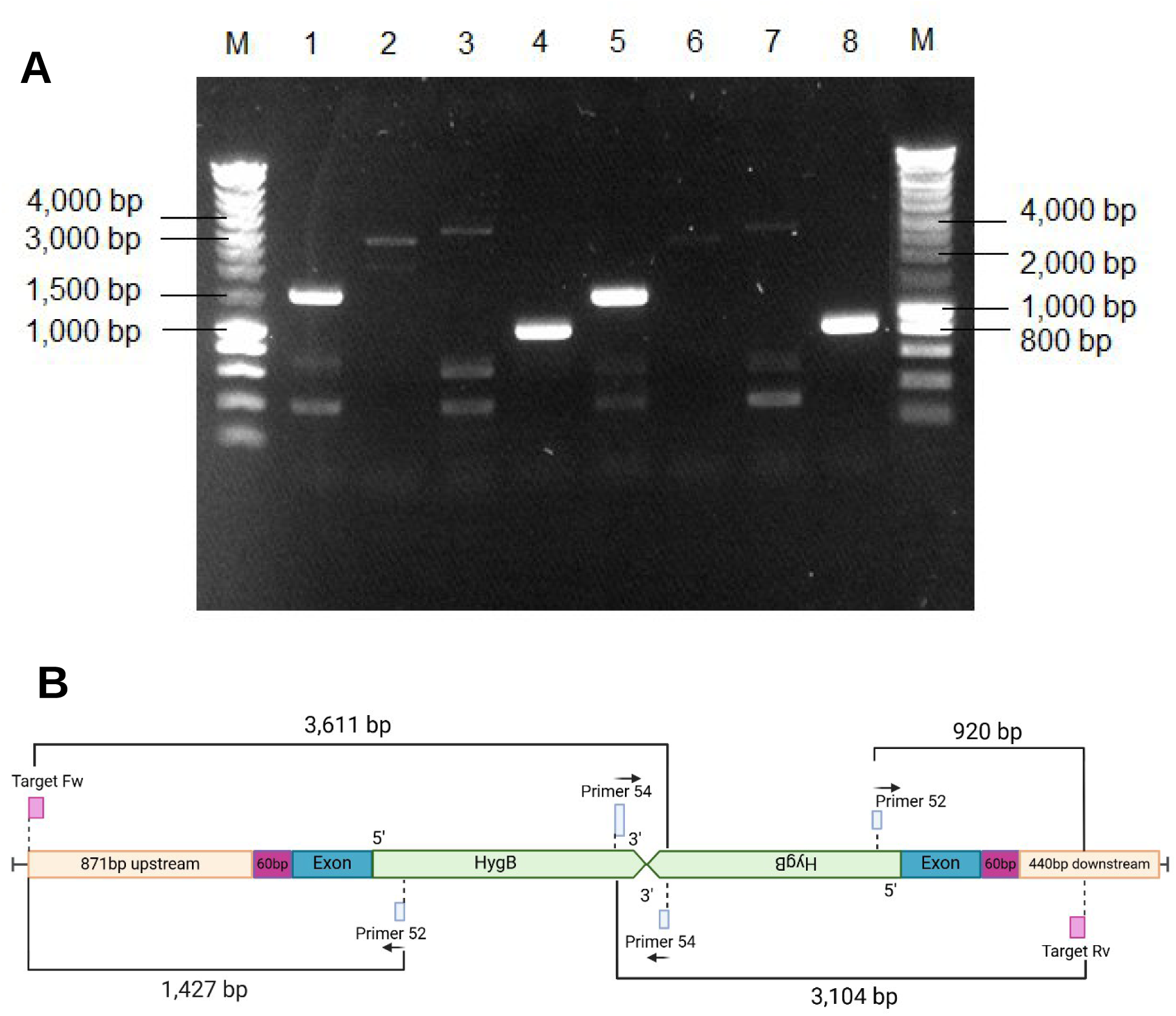
PCR amplification from a *Z. tritici* mutant and a schematic representation indicating the likely insertion event. A) PCR amplifications using combinations of the primers to amplify the target DNA regions and the internal primers for the *HygB* cassette and two replicate DNA extractions from mutant ΔMycgr3107904_1. M: 10-kb (SmartLadder); 1 – 4, ΔMycgr3107904_1 DNA1; 5 – 8, ΔMycgr3107904_1 DNA2. Primers used for lanes 1 and 5: Mycgr3G107904_target Fw + 52; 2 and 6: 54 + Mycgr3G107904_target Rv; 3 and 7: Mycgr3G107904_target Fw + 54; 4 and 8: 52 + Mycgr3G107904_target Rv. B) Schematic representation of the proposed insertion pattern of the *HygB* cassette in mutant ΔMycgr3107904_1 based on PCR and sequencing results.

Sanger sequencing results (Supplementary data) confirmed the insertion pattern proposed in Figure 10B. Sequencing of PCR fragments from lanes 1 and 5 revealed a match with the region between Mycgr3G107904_target Fw (Target Fw in Figure 10B) and internal primer 52 for the *HygB* cassette, indicating an insertion in the 5′ to 3′ orientation. Similarly, sequencing of products from lanes 4 and 8 matched the region between Primer 52 and Mycgr3G107904_target Rv (Target Rv in Figure 10B), confirming an insertion in the 3’ to 5’ orientation.

Additionally, the ∼3,100-bp fragments in lanes 2 and 6 correspond to the expected size of an amplicon between Primer 54 (within the cassette inserted in the 5′ to 3′ orientation) and Mycgr3G107904_target Rv (Target Rv) (Figure 10B). Likewise, the ∼3,600-bp fragments in lanes 3 and 7 match the calculated size of amplification between Primer 54 (from a cassette inserted in the 3′ to 5′ orientation) and Mycgr3G107904_target Fw (Target Fw), further supporting the presence of insertions in both orientations (Figure 10B). Altogether, we genotypically confirmed disruption of the target gene for two mutants of Mycgr3107904.

### Effects on virulence of the *Z. tritici* Mycgr3107904 CRISPR mutants in pathogenicity assays

The first visible disease symptoms appeared at 12 days post inoculation exclusively in plants infected with the wild-type strain, manifesting as small chlorotic lesions. By 14 DPI, these lesions progressed to necrotic tissue, with some showing pycnidia formation on the surface (Figure 11A). In contrast, plants inoculated with either mutant strain, as well as the uninoculated control plants, showed no visible signs of infection at 14 DPI (Figure 11 A, C).

**Figure 11.**
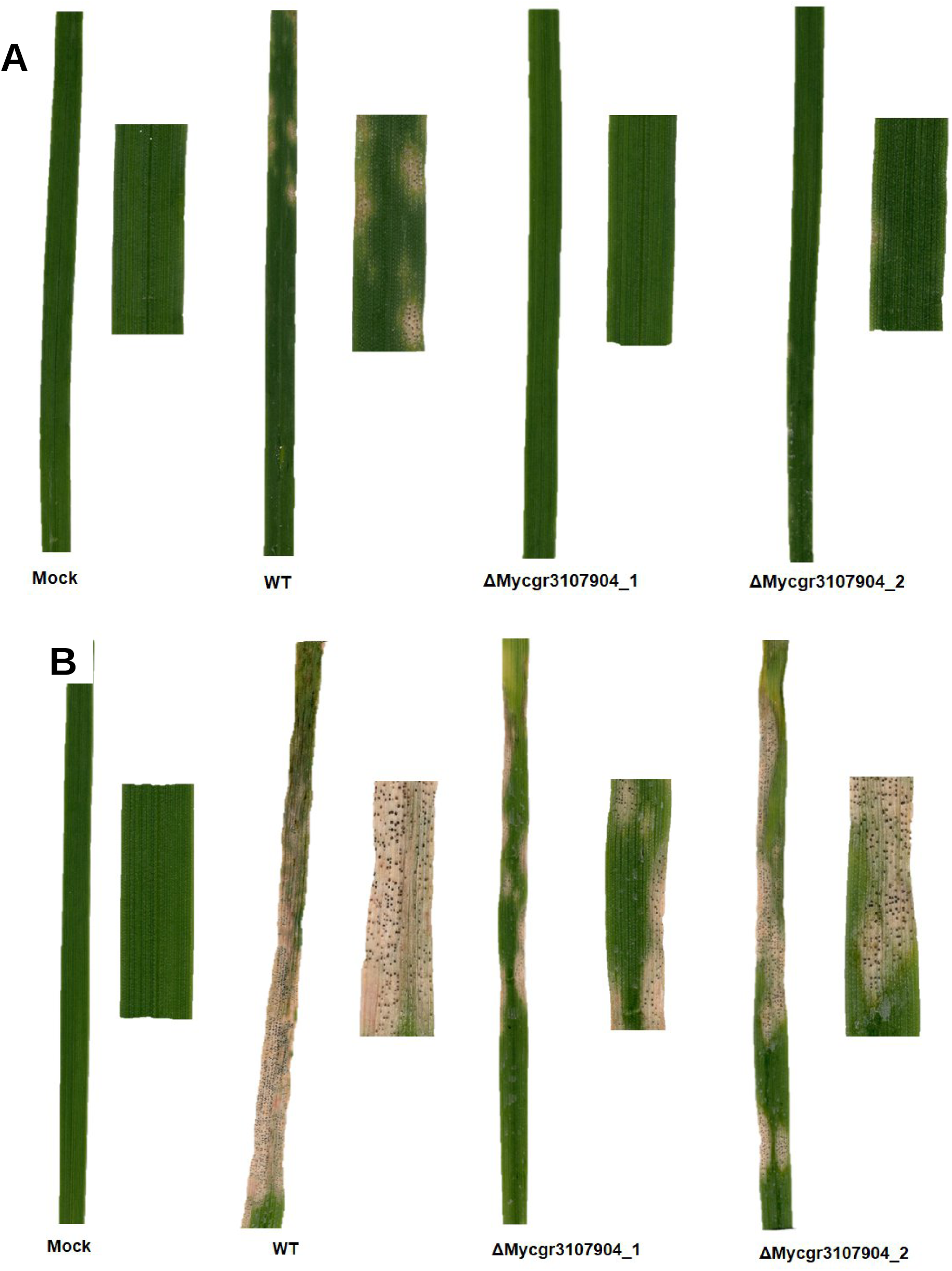
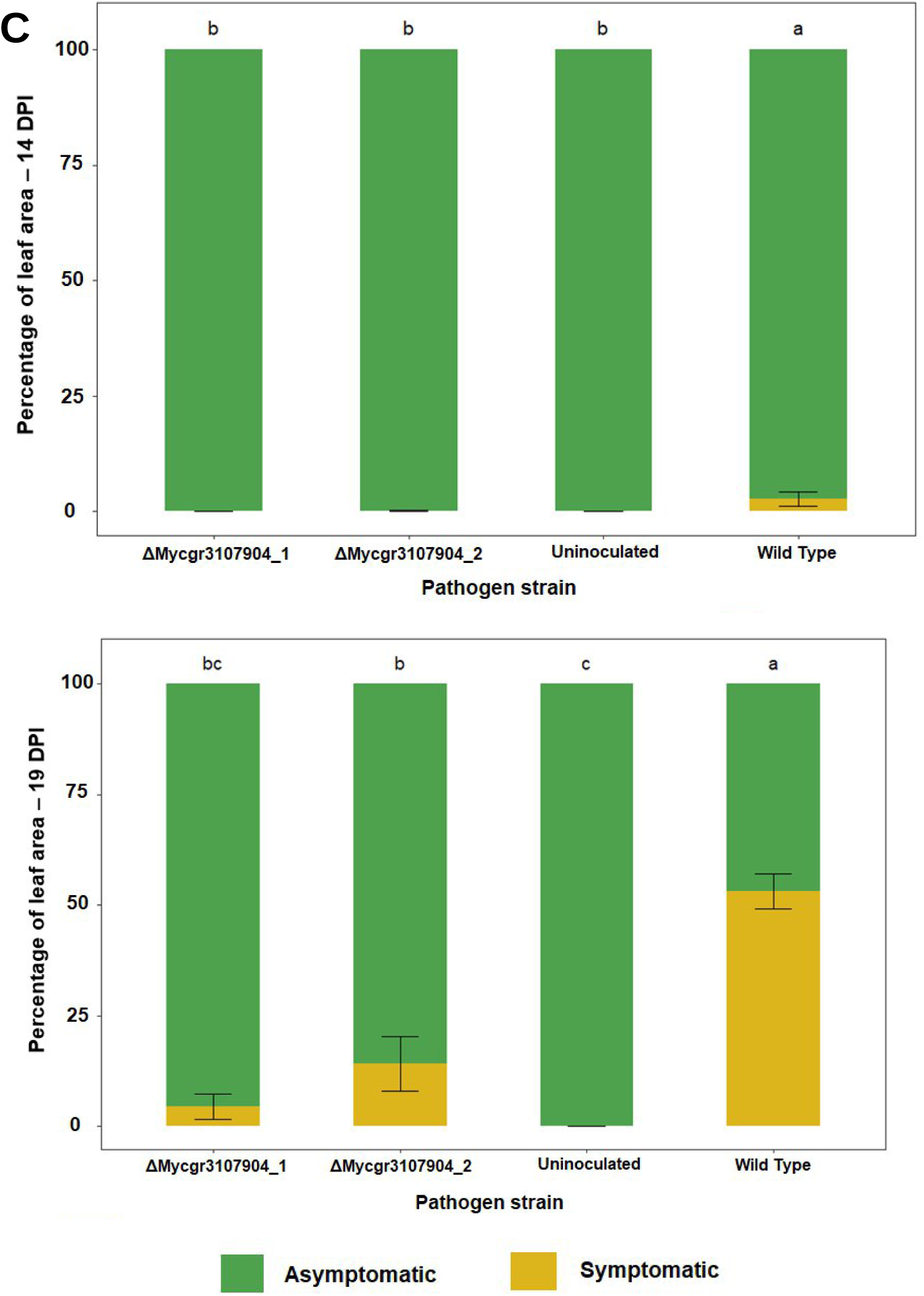
Disease development of the Δ*Mycgr3107904_1* and Δ*Mycgr3107904_2* mutants of *Zymoseptoria tritici* on the susceptible wheat cultivar Taichung 29 compared to wild-type strain IPO323 and mock-inoculated control at A) 14 DPI and B)19 DPI. C) Percentage of symptomatic leaf area of wheat cultivar Taichung29 infected with the Δ*Mycgr3107904_1* or Δ*Mycgr3107904_2* mutants compared to wild type strain IPO323 at 14 and 19 DPI. Stacked bar plots represent the average of one independent replicated experiment with 3 leaves measured at each time point from each inoculation treatment. Error bars are the standard error of the mean. Bars with different letters at the top indicate significant differences (Tukey’s HSD test, P < 0.05).

The Δ*Mycgr3107904_1* and Δ*Mycgr3107904_2* mutants followed the same disease progression steps as IPO323, but with a significant delay. Leaves of Taichung29 inoculated with the mutant strains first developed chlorotic lesions around 17 DPI (based on visual observation). By 19 DPI, these lesions had transitioned to necrosis, following the same two-day interval observed in plants infected with IPO323. However, at this stage, more than 50% of the leaf area in IPO323-inoculated plants was already covered with necrotic lesions and pycnidia (Figure 11 B,C).

Overall, symptom onset and disease progression in the mutants were delayed by approximately 4 to 5 days relative to IPO323. By 21 DPI, leaves infected with IPO323 were completely necrotic and densely covered with pycnidia, reaching the final stage of infection before those inoculated with the mutant strains. Uninoculated control plants showed no signs of disease throughout the experiment.

## Discussion

These results showed clearly that CRISPR/Cas9 with 60-bp flanking sequences could successfully knock out genes in the wheat pathogen *Z. tritici.* Previous attempts by others to use this technique with this pathogen were unsuccessful [36], so our results represent a major advance, even if the number of transformants and the frequency of insertion of the donor DNA into the Cas9 cleavage site were relatively low. The statistically significant reductions in virulence observed in two mutants of one of the targeted effectors proved that this technique can be used for functional analysis of genes in *Z. tritici*.

While in some fungal species like *Fusarium oxysporum* f. sp. lycopersici, longer homology arms actually provide higher transformation efficiencies [53], one key to success in our approach may be that we adapted CRISPR/Cas9 technology to disrupt candidate effector genes in *Z. tritici* using short donor homology flanks (60 bp) to facilitate microhomology-directed repair (MDR) for DSB repair, a strategy that has proven effective in other fungal species [24,32,34]. The advantage of CRISPR/Cas9 is that dsDNA repair through homology can also be achieved using short donor template homology flanks rather than long (>300 bp) homologous regions [32]. For example, in *F. proliferatum*, MDR successfully repaired the DSBs using 50-bp homology regions, while in *B. sorokiniana*, 60-bp flanking regions improved gene knock-out efficiency and integration of the donor template. These approaches allowed for a detailed functional analysis of *FUM1*, a key gene in the fumonisin biosynthetic cluster of *F. proliferatum* [24], and *PKS1*, which encodes a polyketide synthase essential for melanin biosynthesis in *B. sorokiniana* [34]. In *B. cinerea*, Leisen et al. [49]demonstrated that the efficiency of gene targeting increased with donor DNA containing homology flanks up to 60 bp adjacent or distant to the PAM sequence. Their study showed precise insertion of a *FenR* cassette in the correct orientation relative to the target gene (*Bos1*), confirming that short homology flanks can enable efficient gene replacement [32]. Similarly, in our analyses, sequencing of the insertion borders in *Z. tritici* confirmed the successful use of 60-bp homology flanks for insertion of the *HygB* cassette and donor DNA-mediated repair. However, we observed a double insertion of the *HygB* cassette in the Δ*Mycgr3107904_1* mutant, with insertions occurring in both the forward and reverse orientations.

The occurrence of multiple donor insertions in different orientations suggests potential challenges in precisely controlling repair outcomes. Recent work in filamentous fungi has demonstrated that CRISPR/Cas9-induced DSBs can be repaired through the simultaneous action of multiple DNA repair pathways, resulting in complex integration outcomes rather than precise homologous recombination events [28,54]. In *Aspergillus niger*, for example, a protocol for CRISPR/Cas9-mediated targeting of the *PyrG* locus employed a plasmid-based selectable repair cassette designed to be released by cleavage with the same sgRNA/Cas9 complex. This approach revealed that one end of the Cas9-induced DSB was consistently repaired through HDR, whereas the opposing end was frequently resolved by NHEJ [28]. This asymmetric repair resulted in truncated donor integration and retention of sequence signatures corresponding to the original cleavage site. Importantly, this mixed repair outcome occurred despite the presence of homologous sequences, indicating that homologous recombination and NHEJ can act simultaneously on different ends of the same DSB.

Although the donor DNA used in our research was provided as a linear, double-stranded fragment containing short (60-bp) homology arms, the tandem inverted and partially truncated integration of the *HygB* resistance cassette that we observed at the target locus of the Δ*Mycgr3107904_1* mutant is consistent with similar mixed HDR–NHEJ repair processes. In *Z. tritici,* as previously reported in *A. niger,* DNA repair was likely initiated by homologous recombination (HR) on one side of the break, sufficient to anchor the donor DNA and confer selection, while the second end of the break remains permissive to NHEJ-mediated end joining. Under these conditions, linear donor fragments can be captured in multiple copies and ligated in either orientation, with end trimming prior to ligation resulting in partial loss of donor sequences.

Together, these observations support a model in which CRISPR/Cas9-induced DSBs in *Z. tritici* likely give rise to asymmetric repair outcomes that favor complex donor integration, particularly when NHEJ remains active.

In *Magnaporthe oryzae,* another fungal model system, the repair pathways of Cas12a- and Cas9-induced DSBs resulted in a spectrum of DNA mutations. Genotyping of multiple transformants revealed insertions and deletions (INDELs), deletion plus insertion events, simple donor DNA insertions, large concatemer insertions and genomic deletions, some producing drastic mutations at the targeted loci [55]. Notably, integration of the hygromycin resistance cassette at the target locus was unsuccessful in a *M. oryzae* strain lacking *Ku80*, a key component of the NHEJ pathway, despite frequent integration under identical conditions in the wild-type background. This observation suggests that *Ku80*-dependent NHEJ is required for efficient donor DNA integration in this system [55]. These findings indicate that repair outcomes following CRISPR/Cas12a- or Cas9-induced DSBs in fungi depend, at least in part, on error-prone end-joining mechanisms. Additionally, PCR genotyping of *M. oryzae* transformants and long-read sequencing of amplified products at the target locus revealed that they harbored an almost complete hygromycin-coding sequence plus an additional 140-bp hygromycin fragment. The inserted donor DNA was frequently fragmented and rearranged, with hygromycin sequences detected in both forward and reverse orientations [55]. These findings align with the tandem and inverted insertions of the *HygB* cassette observed at the target locus of *Mycgr3107904,* underscoring the structural complexity of CRISPR-mediated repair outcomes possibly associated with the involvement of the NHEJ pathway for donor DNA integration.

In the *ΔMycgr3107904_2* mutant, sequencing revealed the insertion of a nucleotide sequence (∼298 bp) that was unrelated to the *HygB* cassette, suggesting that NHEJ may also have been the repair mechanism activated instead of MDR. In fact, given that NHEJ is often the predominant repair mechanism in fungi [21], it is likely that error-prone NHEJ also contributed to the observed repair events in the *ΔMycgr3107904_2* mutant. While NHEJ has traditionally been associated with small INDELs of less than 50 bp at Cas9-sgRNA cleavage sites, recent studies indicate that CRISPR/Cas9 can also lead to unintended, large genomic modifications at target sites [56,57]. These include large insertions [58] similar to the long fragment that was inserted in the *ΔMycgr3107904_2* mutant. To fully characterize both the long insertion of random DNA in *ΔMycgr3107904_2* and the tandem and inverted insertion of the *HygB* cassette in *ΔMycgr3107904_*1, long-range PCR followed by long-read sequencing will be required, as this approach would allow unambiguous determination of the full length, structure, and sequence identity of the inserted DNAs beyond what can be resolved from overlapping Sanger sequencing reads.

Antibiotic-resistant colonies were obtained for Mycgr3109710 and Mycgr394290, but PCR and sequencing analyses revealed undisrupted target genes, indicating failed target-site integration. Antibiotic resistance alone does not confirm cassette insertion at the designed target site; several studies report resistant transformants lacking the selectable marker at the intended locus, attributed to ectopic integration or Cas9 off-target activity [59–61]. For example, CRISPR/Cas9 editing in *Candida auris* generated 4,900 drug-resistant transformants in which PCR analysis failed to detect correct cassette integration at the target gene, indicating that resistance was conferred by integration at secondary genomic sites [60]. Cas9 can cleave partially complementary sequences, particularly tolerating mismatches outside the PAM-proximal seed region where guide–DNA structural flexibility permits cleavage [62,63] During transformation, exogenous donor DNA can integrate at these unintended DSBs through error-prone end-joining pathways [64] We conclude that the antibiotic resistance observed in the *ΔMycgr3109710* and *ΔMycgr394290 Z. tritici* strains arose from ectopic insertion of the resistance cassette rather than precise on-target integration.

The relatively low frequency of transformed colonies for Mycgr3107904, Mycgr3109710, and Mycgr394290, combined with the complete absence of colonies for Mycgr3106502 and Mycgr3103393, suggests locus-dependent variation in DNA repair pathway usage. Such variation has been documented in other organisms, where NHEJ, HR, and MDR occur at unequal frequencies across the genome [54,65]. Local genomic context including flanking repetitive DNA sequences, chromatin organization, chromosome location, and regional sequence composition can influence which repair pathways are preferentially activated [54]. We hypothesize that these factors collectively establish a locus-specific repair pathway preference in *Z. tritici* that determines the efficiency and outcomes of Cas9-mediated genome editing.

Previously, Khan et al. [36] attempted to implement CRISPR/Cas9-mediated transformation in *Z. tritici* by targeting the *PyrG* gene, which is not required for pathogenicity and allows growth on 5’-fluorotic acid selection media when mutated. Initially, a *Z. tritici* strain expressing Cas9 was generated, and protoplasts were transformed with sgRNA using PEG-mediated transformation. Although 14 colonies grew on selection media, sequencing revealed no mutations in *PyrG* [47]. In contrast, we did not express Cas9 in *Z. tritici* either through a plasmid vector or by generating a Cas9-expressing strain. Our approach may have helped prevent Cas9 degradation, which Khan et al. [36] suggested as a possible issue in their study. We employed direct transformation of fungal protoplasts via PEG-mediated uptake of the RNP complex, avoiding the need for fungal expression of Cas9, which has been associated with unintended toxic effects in *M. oryzae* and other fungi [27,66].

Khan et al. [36] also tested whether the lack of mutations was gene specific by targeting a different gene, *NiaD* (nitrate reductase), using an RNP complex along with a donor template with 1-kb homologous regions flanking the *NiaD* cleavage site for HDR. Despite obtaining 17 colonies on selection media, none showed successful integration of the donor cassette [36]. In contrast, our approach used a donor template with short (60 bp) flanking regions homologous to the upstream and downstream sequences of the target gene, rather than the sgRNA cleavage site. This key difference from previous attempts may have played a role in successfully disrupting the target gene. The use of short homologous flanks of only 60 bp, which has proved to be effective in other fungi, might have helped to minimize the risk of altering adjacent gene sequences [24], and in our case, it enabled successful transformation of one of the target effector genes, with integration of the *HygB* cassette at the intended site. In the current study, we demonstrated that 60-bp flanking regions are sufficient for disruption of a targeted genomic locus in *Z. tritici*.

Despite successfully generating transformants, our approach faced several challenges, including the extremely low number of colonies obtained—only one to two colonies on selective media for three target genes and none for two of the five effector genes. Our protoplast regeneration strategy followed a method commonly used for filamentous fungi including *Fusarium* and *Pseudocercospora fijiensis*, where transformed protoplasts were embedded in regeneration medium (RM) without antibiotics for 18 hours before overlaying the plates with RM containing hygromycin. However, since *Z. tritici* predominantly grows in a yeast-like form rather than forming hyphae in *in-vitro* culture, this approach likely hindered the emergence of transformed colonies to the surface of the medium. To improve colony recovery, a modification in the post-transformation culture method by first growing protoplasts in liquid RM before plating them onto solid RM with hygromycin may be required. Additionally, we aim to optimize the protoplast generation process to increase yield, as our current method barely reached the minimum protoplast concentration reported in other fungal transformation protocols, such as those for *P. fijiensis.¶*

Regardless of the challenges encountered, our results demonstrate the feasibility of using CRISPR/Cas9 for gene knockouts in *Z. tritici* and highlight the need for further optimization. Refining conditions such as donor DNA concentration, transiently limiting Cas9 activity, or biasing repair toward HDR could enhance the precision of genome editing in *Z. tritici*. A key advantage of CRISPR/Cas9 is that, once implemented in a given organism, it allows for flexible genome editing by simply modifying the sgRNA protospacer sequence. Additionally, multiple genes can be targeted simultaneously by introducing multiple sgRNAs along with Cas9 [67], enabling efficient generation of mutant strains for functional studies. This is particularly valuable for *Z. tritici*, where effector gene redundancy has been observed [20], and multi-gene knockouts could help clarify the roles of virulence-related genes.

There are no previous reports of transformants in *Z. tritici* obtained through implementation of CRISPR/Cas9 technology, and *Mycgr3107904* has not been disrupted previously. However, successful knockouts of Hce-2-domain-containing effector genes, to which Mycgr3107904 belongs, have been reported in other fungi. In the apple pathogen *Valsa mali*, the double-knockout strain *ΔVmHEP&VmHEP2* showed no significant differences in hyphal growth compared to the wild-type strain when grown on PDA. This contrasts with our findings, where *ΔMycgr3107904_1* exhibited delayed melanization, a faster growth rate on YSA, YMA, and PDA, and more than doubled the colony diameter of the wild-type strain on PDA. Additionally, growth on water agar (WA) revealed that while the wild-type strain developed hyphal growth, mutant *ΔMycgr3107904_1* showed almost none after 14 days and mutant *ΔMycgr3107904_2* also showed significantly less hyphal growth at 14 days. The observed differences in colony morphology, melanization, and growth rate of mutant *ΔMycgr3107904_1* are likely caused by off-target disruptions rather than disruption of the target gene. Although we confirmed disruption of the candidate effector sequence, the alteration of a single predicted effector is unlikely to produce such a pronounced phenotypic change. Supporting evidence comes from mutant *ΔMycgr3107904_2*, which harbors the same target gene disruption but lacks the altered phenotype observed in *ΔMycgr3107904_1*. To conclusively determine whether Mycgr3107904 affects growth and other physiological processes in *Z. tritici*, additional independent mutant colonies must be generated and phenotypically characterized. Complete genomic sequencing of the *ΔMycgr3107904_1* mutant would identify which other genes may have been altered to give the observed phenotypes.¶

Unintended phenotypes following CRISPR/Cas9-mediated genome editing are relatively common. In *A. niger,* 24% of CRISPR/Cas9-generated mutants lacking the *xlnR* gene, which encodes a (hemi-)cellulolytic transcriptional regulator, and derived from wild-type strains exhibited aberrant growth phenotypes. In contrast, no such phenotypes were observed in a NHEJ-deficient *ΔxlnR* mutant, suggesting that the unexpected phenotypes were caused by CRISPR/Cas9-induced off-target mutations rather than *xlnR* disruption [64]. Whole-genome sequencing further showed that mutants generated in the NHEJ-deficient background accumulated significantly fewer mutations, consistent with greater genomic instability in wild-type strains where both the NHEJ and HDR repair pathways are active [64]. Similarly, the altered colony morphology, growth rate, and melanization observed in the *ΔMycgr3107904_1* mutant of *Z. tritici* may reflect additional mutations acquired during transformation or off-target effects, which could be resolved through genome-wide analysis.

In *V. mali*, the double mutant *ΔVmHEP&VmHEP2* showed reduced virulence on apple leaves, while single mutants were not altered in virulence. In *Verticillium dahliae*, the Hce2-domain-containing effector *VdCE11* was knocked out, and the mutant strain was used to inoculate cotton and *Arabidopsis* plants. In both hosts, disease development was weakened, and virulence was restored upon complementation, confirming the role of *VdCE11* in pathogenicity [68]. In our analyses, we disrupted *Mycgr3107904*, one of three Hce2-domain-containing effectors in *Z. tritici*, and showed that the knockout mutant of this gene had significantly delayed disease progression in the susceptible wheat cultivar Taichung 29. Further optimization of CRISPR/Cas9 in *Z. tritici* and the generation of double and triple mutants will help determine whether the other two Hce2-domain-containing effectors also contribute to virulence.

The observed differences in virulence are particularly intriguing because early studies described the Hce2-domain-containing effector ZtNIP1 from *Z. tritici* as a necrosis-inducing protein in wheat when expressed in *Pichia pastoris,* supporting a direct role in virulence [6]. However, subsequent work reclassified this protein, now termed Zt-KP4-1, as a structural homolog of fungal killer proteins, demonstrating antifungal activity but no necrosis induction in wheat [69]. Based on this revised functional annotation, we hypothesize that candidate effector *Mycgr3107904* has an indirect, fitness-related role rather a classical host-manipulating function. Rather than triggering host cell death, *Mycgr3107904* may enhance pathogen success by suppressing competing fungi or regulating microbial interactions within infected tissue. The altered virulence phenotypes observed for the two mutants of *Mycgr3107904* may therefore arise from reduced competitive fitness or impaired colonization dynamics during infection, highlighting a possible role for killer protein-like effectors in shaping the infection environment rather than directly manipulating host immunity. Further research is needed to elucidate the specific function of this effector and the development of a complementation mutant, which would confirm the involvement of *Mycgr3107904* in the virulence of *Z. tritici* to wheat.

## Acknowledgements

The authors thank Els Verstappen for technical support.

## Funding

This research was funded by the United States Department of Agriculture, Agricultural Research Service (USDA-ARS) research project 5020-21220-014-00D.

## Notes

### Competing Interest Statement

The authors have declared no competing interest.

